# Itaconate and derivatives reduce interferon responses and inflammation in influenza A virus infection

**DOI:** 10.1101/2021.01.20.427392

**Authors:** Aaqib Sohail, Azeem A. Iqbal, Nishika Sahini, Mohamed Tantawy, Moritz Winterhoff, Thomas Ebensen, Robert Geffers, Klaus Schughart, Fangfang Chen, Matthias Preusse, Marina C. Pils, Carlos A. Guzman, Ahmed Mostafa, Stephan Pleschka, Christine Falk, Alessandro Michelucci, Frank Pessler

**Author notes:** Joint first authors (Aaqib Sohail, Azeem A. Iqbal, Nishika Sahini). Corresponding author: Frank Pessler, MD, PhD.

## Abstract

Itaconate has recently emerged as a metabolite with immunomodulatory properties. We evaluated effects of endogenous itaconate and exogenous itaconate, dimethyl-, and 4-octyl-itaconate on host responses to influenza A virus infection. Infection induced *ACOD1* (the enzyme catalyzing itaconate synthesis) mRNA in monocytes and macrophages, which correlated with viral replication and was abrogated by itaconate treatment. Pulmonary inflammation and weight loss were greater in *Acod1^-/-^* than wild-type mice, and ectopic synthesis of itaconate in human epithelial cells reduced infection-induced inflammation. The compounds induced different recruitment programs in infected human macrophages, and transcriptome profiling revealed that they reversed infection-triggered interferon responses and modulated inflammation in cell lines, PBMC, and lung tissue. Single-cell RNA sequencing of PBMC revealed that infection induced *ACOD1* exclusively in monocytes, whereas treatment silenced IFN-responses in monocytes, lymphocytes, and NK cells. Viral replication did not increase under treatment despite the dramatically repressed IFN responses, but 4-octyl itaconate inhibited viral transcription in PBMC. The results reveal dramatic reprogramming of host responses by itaconate and derivatives and their potential as adjunct treatments for hyperinflammation in viral infection.

## Introduction

As exemplified by influenza viruses (Ramos & Fernandez-Sesma, 2015) and pathogenic coronaviruses (Ragab, Salah Eldin et al., 2020), overshooting host inflammation, oxidative stress and cell damage/death are major determinants of morbidity and mortality in severe viral infections of humans. Consequently, there is a great need for adjunct treatments that can be given in conjunction with direct-acting anti-virals, with the aim to modulate the host response in such a way that end-organ damage is reduced and clinical outcome is improved.

Itaconic acid initially attracted great interest in the field of biotechnology due to its highly reactive double bonds, which make it an efficient intermediate in the biosynthesis of industrial polymers (reviewed in (Cordes, Michelucci et al., 2015)). The discoveries that its synthesis is highly induced during macrophage activation (Strelko, Lu et al., 2011) and that *immune response gene 1* (*Irg1,* since then renamed *aconitate decarboxylase*) encodes the enzyme cis-aconate decarboxylase (ACOD1 in humans, CAD in other organisms), which converts cis-aconitate to itaconate (Chen, Lukat et al., 2019, Michelucci, Cordes et al., 2013), have triggered intense efforts to elucidate functions of the endogenous ACOD1/itaconate axis in inflammation and infection. The early recognition that itaconate possesses anti-inflammatory and immunomodulatory properties also suggested that itaconate and prodrugs derived from it may have potential as immunomodulatory and cytoprotective interventions for inflammation-driven diseases (reviewed in (Hooftman & O’Neill, 2019, O’Neill & Artyomov, 2019)).

Most evidence of anti-inflammatory and immunomodulatory effects of itaconate has actually been obtained by treating cells or small animals with chemically modified variants that are presumably taken up more efficiently. 4-Octyl-itaconate (4OI) contains an added 8-carbon chain and acts primarily through activation of the NRF2 pathway (Mills, Ryan et al., 2018). It has, for instance, shown cytoprotective and anti-inflammatory effects in a variety of models, including lipopolysaccharide (LPS)-induced macrophage activation and LPS-induced sepsis in mice (Mills et al., 2018), in a neuronal model of oxidative stress (Liu, Feng et al., 2018), and in peripheral blood mononuclear cells (PBMC) from patients with systemic lupus erythematosus (Tang, Wang et al., 2018). Dimethyl-itaconate (DI) has also been shown to exert NRF2 independent anti-inflammatory effects by inhibiting IκBζ activity (Bambouskova, Gorvel et al., 2018). Furthermore, DI possesses cytoprotective effects, which was, for instance, shown by reduced cell damage in mouse models of cardiac ischemia (Lampropoulou, Sergushichev et al., 2016).

However, cytoprotective properties have also been demonstrated for unmodified itaconate, notably in a rat model of cerebral ischemic-reperfusion injury (Cordes, Lucas et al., 2020). Both compounds likely exert cytoprotective effects by reducing reactive oxygen species (ROS) generation via inhibition of succinate dehydrogenase (Cordes et al., 2020, Cordes, Wallace et al., 2016, Lampropoulou et al., 2016). Considering these varied effects of itaconate and derivatives, it is not surprising that administration of DI has demonstrated beneficial effects in a variety of mouse models of infection-associated inflammation such as LPS-induced mastitis (Zhao, Jiang et al., 2019) and endometritis (Xu, Jiang et al., 2020b), and fungal keratitis (Gu, Lin et al., 2020). Following early evidence that *Acod1* transcription is strongly induced in severe influenza A virus (IAV) infection in mice (Preusse, Tantawy et al., 2013), its impact on host susceptibility to viral infections has also been studied, and protective effects of itaconate on neurons exposed to RNA viruses have been reported (Cho, Proll et al., 2013, Daniels, Kofman et al., 2019, Passalacqua, Purdy et al., 2019). On the other hand, in a recent study using a mouse model of IAV infection, deleting the *Acod1* locus did not affect weight loss or survival, whereas it clearly worsened the outcome of *Mycobacterium tuberculosis* infection in the same mouse strain (Nair, Huynh et al., 2018). This lack of effect on IAV infection was surprising because abnormally increased inflammation is usually asociated with deleterious effects on the host during viral infections. For instance, it has been shown that overactive inflammatory responses are associated with decreased survival and increased weight loss in IAV infected inbred mouse strains (Alberts, Srivastava et al., 2010). While the role of endogenous itaconate in protection from viral infections thus remains incompletely understood, the above results do suggest a strong potential of external application of itaconate or its derivatives to ameliorate inflammation and other deleterious consequences of viral infections.

Focusing on IAV as one of the most important human respiratory viral pathogens (Collaborators, 2018), we therefore examined role and regulation of the endogenous ACOD1/itaconate axis in a mouse model of IAV infection and subsequently performed a detailed analysis of the modulatory effects of exogenous addition of itaconate and derivatives on inflammatory responses in IAV-infected human cell lines, explanted lung tissue, and PBMC, both at the bulk and single-cell levels. We find that loss of ACOD1 and itaconate synthesis led to increased inflammation during infection and that exogenous itaconate, DI, and 4OI substantially reduced inflammation, most notably IFN responses, without enhancing viral replication. 4OI was unique in that it also exerted a direct anti-viral effect.

## Results

### *Acod1* expression and itaconate synthesis are induced in mouse lung during IAV-infection and limit pulmonary inflammation and disease severity

Compared with C57BL/6J, DBA/2J mice are more susceptible to IAV infection in that weight loss is more rapid, mortality is higher, and inflammation is more pronounced (Preusse, Schughart et al., 2017, Srivastava, Blazejewska et al., 2009). We have shown previously that *Acod1* transcription in the first 48 h of infection is higher in IAV-infected lung from DBA/2J than from C57BL/6J mice (Preusse et al., 2013). A reanalysis of our RNAseq data from a longer time course (Wilk, Pandey et al., 2015) showed that this difference persisted at least through day 3 post infection (p.i.) and that *Acod1* expression nearly normalized by day 14 in the surviving strain (Fig. 1A). This analysis also revealed a remarkable correlation with expression of *Tnfaip3* (encoding an anti-inflammatory deubiquitinase) and a weaker positive correlation with *Hmox1* and *2* (encoding inducible anti-oxidant enzymes), all of which are known to be co-regulated with *Acod1* (Li, Zhang et al., 2013, Uddin, Joe et al., 2016). *Acod1* was the 4^th^ most highly upregulated pulmonary mRNA 48 h p.i. in C57BL/6J mice, where it appeared in a cluster of transcripts pertaining to IFN responses and innate immunity (Fig. 1B). These results underscored the close association of *Acod1* expression with systemic inflammation and oxidative stress in IAV-infected mice. We therefore tested whether targeted deletion of *Acod1* would affect inflammatory responses and host susceptibility to IAV infection. Itaconate was not detectable in lungs from IAV-infected *Acod1*^-/-^ mice; it was highly elevated on days 7/8 p.i. in the *Acod1^+/+^* and, significantly less, in *Acod1^+/-^* mice, and returned to normal by day 14 (Fig. 1C). Both weight loss and mortality were higher in *Acod1^-/-^* than *Acod^+/+^* female C57BL/6J mice (Fig. 1D,E), with greatest changes occurring around day 8, the time of maximal itaconate accumulation. Histologically assessed pulmonary inflammation was greatest in *Acod1*^-/-^ mice and intermediate in *Acod1*^+/-^ mice (Fig. 1F). Thus, itaconate synthesis correlated with disease activity, and loss of *Acod1* was associated with more severe disease and higher pulmonary inflammation.

**Figure 1.**
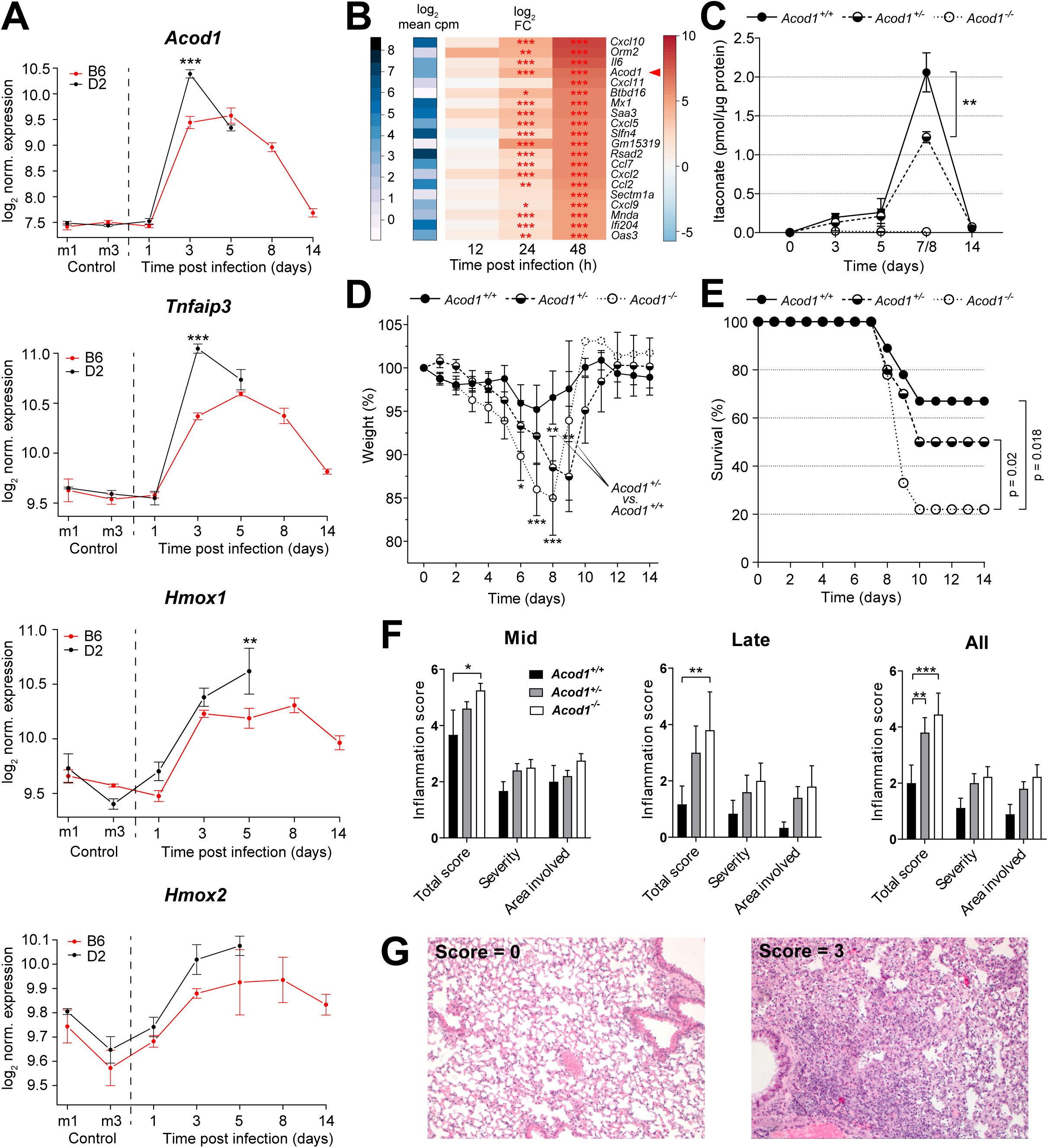
Reciprocal interactions between *Acod1* expression and inflammation during influenza A virus infection in mice. **A.** Higher pulmonary expression of *Acod1, Tnfaip3,* and *Hmox1* and *2* mRNA in DBA/2J (D2) than in C57BL/6J (B6) mice following IAV infection (reanalysis of published RNAseq data (Wilk et al., 2015)) (n=3 each strain and time point). *p<0.05; **p<0.01; ***p<0.001 (pairwise t-test with Bonferroni-Holm multiple comparisons correction). **B.** *Acod1* mRNA is among the most highly upregulated transcripts in mouse lung within 48 h of IAV infection (RNAseq, n=3 per time point). **C.** Lower pulmonary levels of itaconate in IAV-infected *Acod1^+/-^* than in *Acod1^+/+^* mice. Itaconate was not detected in the *Acod1^-/-^* mice (n=36 *Acod1*^+/+^, 27 *Acod1*^+/-^, 4 *Acod1*^-/-^ mice). *p<0.05; **p<0.01; ***p<0.001 (unpaired t-test). **D.** Greater weight loss of female *Acod1^-/-^* C57BL/6J mice during infection with IAV. Mice had to be euthanized when weight loss exceeded 20%; n=9 *Acod1^+/+^*, 10 *Acod1^+/-^*, 9 *Acod1^-/-^* mice. *p<0.05; **p<0.01; ***p<0.001 (unpaired t-test) **E.** Survival data of the same experiment as in C. Median survival after day 7 was significantly lower in *Acod1^-/-^* mice (p=0.018, Mann-Whitney-U test). **F.** Semi-quantitative histopathological scoring of lungs on days 8/9 (mid), 11 and 14 (late) reveals highest inflammation in *Acod1^-/-^* mice; n=9 *Acod1^+/+^*, 10 *Acod1^+/-^,* 9 *Acod1^-/-^* mice (all female). *p<0.05; **p<0.01; ***p<0.001 (two-way ANOVA, Tukey’s multiple comparisons test) **G.** H&E stained lung sections showing inflammation severity scores of 0 (left) and 3 (right).

### IAV infection induces *ACOD1* expression in human PBMC and macrophages

We then tested whether IAV infection increased *ACOD1* mRNA levels in human PBMC and M1/M2 macrophages. Phorbol myristate acetate (PMA)-differentiated THP-1 (dTHP-1) cells were analyzed for comparison. Viral transcription was highest in M2 cells (Fig. 2A). The strongest *ACOD1* mRNA induction was seen in M2 cells and undifferentiated PBMC, and the weakest in M1 cells and dTHP-1 cells (Fig. 2B-E), even though the latter two cell types did support a significant degree of viral transcription (Fig. 2A,D). As in IAV-infected mouse lung (Fig. 1), *TNFAIP3* and *ACOD1* transcriptional correlated with each other (Fig. 2C). Infection of dTHP-1 cells with three IAV (H1N1) strains of differential replication efficiency (Petersen, Mostafa et al., 2018b) showed the strongest *ACOD1* and *TNFAIP3* induction by the two strains with the highest and the weakest by the strain with the lowest replication efficiency (Fig. 2D,E). Thus, the extent of *ACOD1* induction after IAV infection differed significantly among the cell types tested, was strongest in M2 primary cells, was accompanied in all cases by an increase in *TNFAIP3* mRNA, and correlated with viral replication.

**Figure 2.**
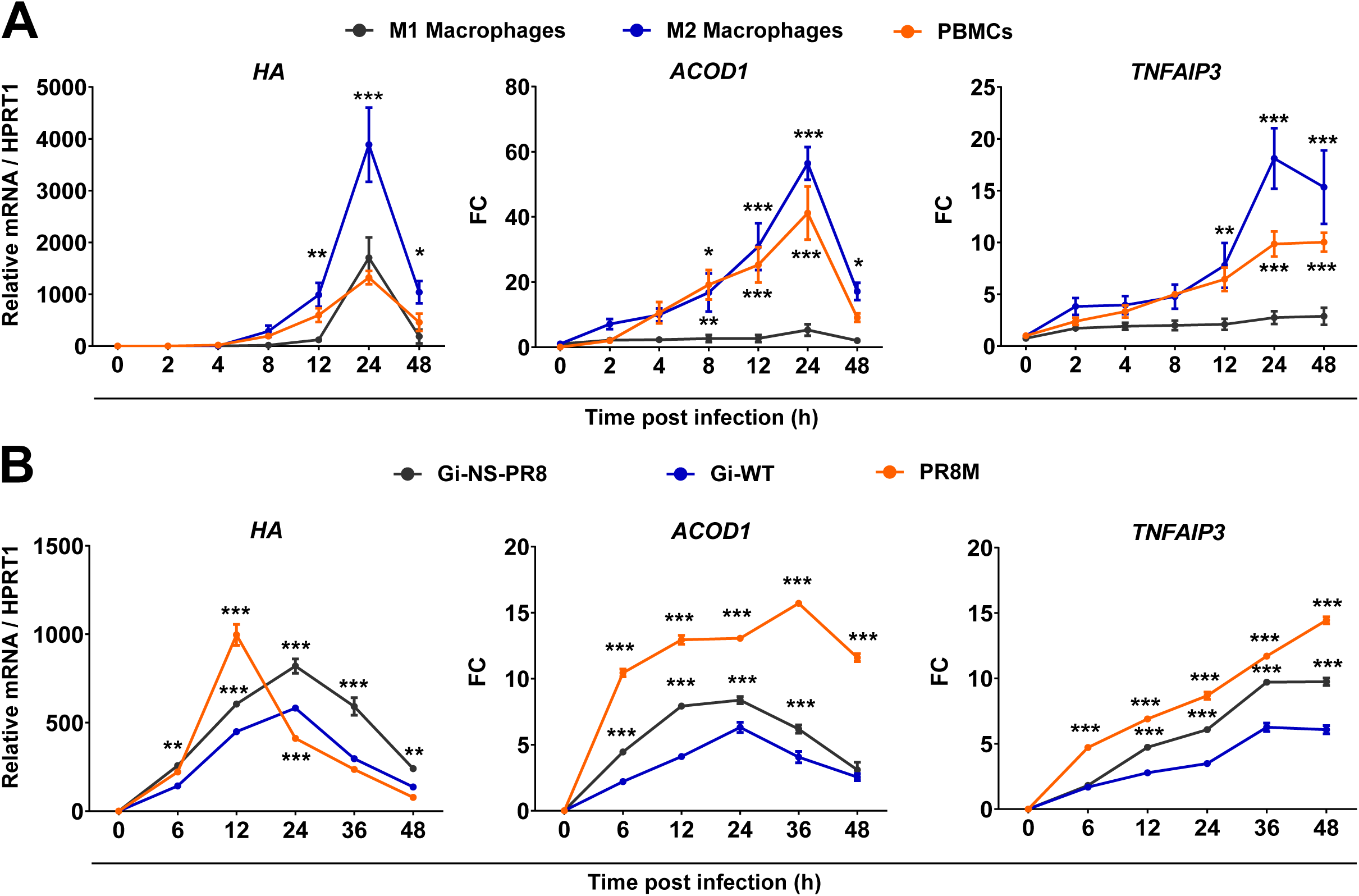
Differences among human myeloid cells in *ACOD1* mRNA induction during IAV infection. **A.** Induction of *ACOD1* and *TNFAIP3* mRNA (RT-qPCR) in M1 and M2 macrophages and PBMCs during IAV infection. **B.** Induction of *ACOD1* and *TNFAIP3* (RT-qPCR) in dTHP-1 cells correlates with virulence of three IAV strains. Mean ±SEM (n=4). *HA* = IAV hemagglutinin. *p<0.05; **p<0.01; ***p<0.001 (two-way ANOVA, Tukey’s multiple comparisons test, with reference to M1 macrophages in A, and to Gi-WT in B).

### Itaconate and DI markedly reduce IFN responses in dTHP-1 cells without affecting viral RNA replication

We then tested whether these compounds could modulate cellular inflammation due to IAV infection. In infected dTHP-1 cells, treatments with itaconate or DI did not affect viral hemagglutinin (*HA)* mRNA expression, but both reduced induction of *CXCL10*, a key pro-inflammatory chemokine during influenza virus infection, with DI being several-fold more potent (Fig. 3B,C, Fig. S1). The 12 h time point was chosen because our previous work had identified it as the peak of IFN-I responses at the mRNA level in IAV-infected dTHP-1 cells (Petersen, Mostafa et al., 2018a). We then assessed global effects of itaconate and DI on cellular gene mRNA expression. A comparison of differentially expressed genes (DEGs) suggested that the two compounds had both shared and unique effects on gene expression (Fig. 3D). Both compounds had considerable effects on gene expression in uninfected cells (Fig. 3E,F). As expected, IAV infection of untreated cells caused a major induction of IFN-regulated transcripts by 12 h p.i. (see mRNAs labeled in the volcano plot in Fig. 3G). Strikingly, addition of both itaconate and DI led to global down-regulation of IFN-regulated genes. This was accompanied by upregulation of a substantial number of other transcripts, indicating that the compounds also modulated other cellular processes (Fig. 3H,I). A principle component analysis (PCA) indicated that effects of the compounds on the cells were so pronounced at these concentrations that treated infected and treated uninfected cells clustered closely together, but far away from untreated infected or uninfected cells (Fig. S2). Indeed, a hierarchical clustering analysis revealed an across-the-board normalization of a clade of IFN- and inflammation-related transcripts by the compounds, but also clusters of DEGs that were upregulated by either or both compounds with respect to both uninfected and infected cells, further indicating effects upon cell homeostasis in general (Fig. S3). mRNA and protein expression may be differentially affected by IAV infection due to virus-induced “host-cell shut-off”. We therefore assessed release of inflammation-related polypeptides in supernatants from the same experiment (Fig. 3J). This analysis confirmed the downregulation of pro-inflammatory factors (e.g., IP-10, MCP-1) by itaconate and DI, but also showed increased IL-8 (DI only) and minor increases of IL1B (both IA and DI) in uninfected cells. Of note, both compounds also increased levels of the anti-inflammatory IL1 receptor antagonist (IL1RA) in supernatants from both infected and uninfected cells. A comprehensive view of the protein targets that were detected by this assay revealed that the two compounds effected substantially different changes in the target cells, as several factors were induced even in uninfected cells specifically by DI or itaconate (Fig. S4). Thus, itaconate and DI apparently act on macrophages to release different recruitment programs that modify local inflammatory cell populations.

**Figure 3.**
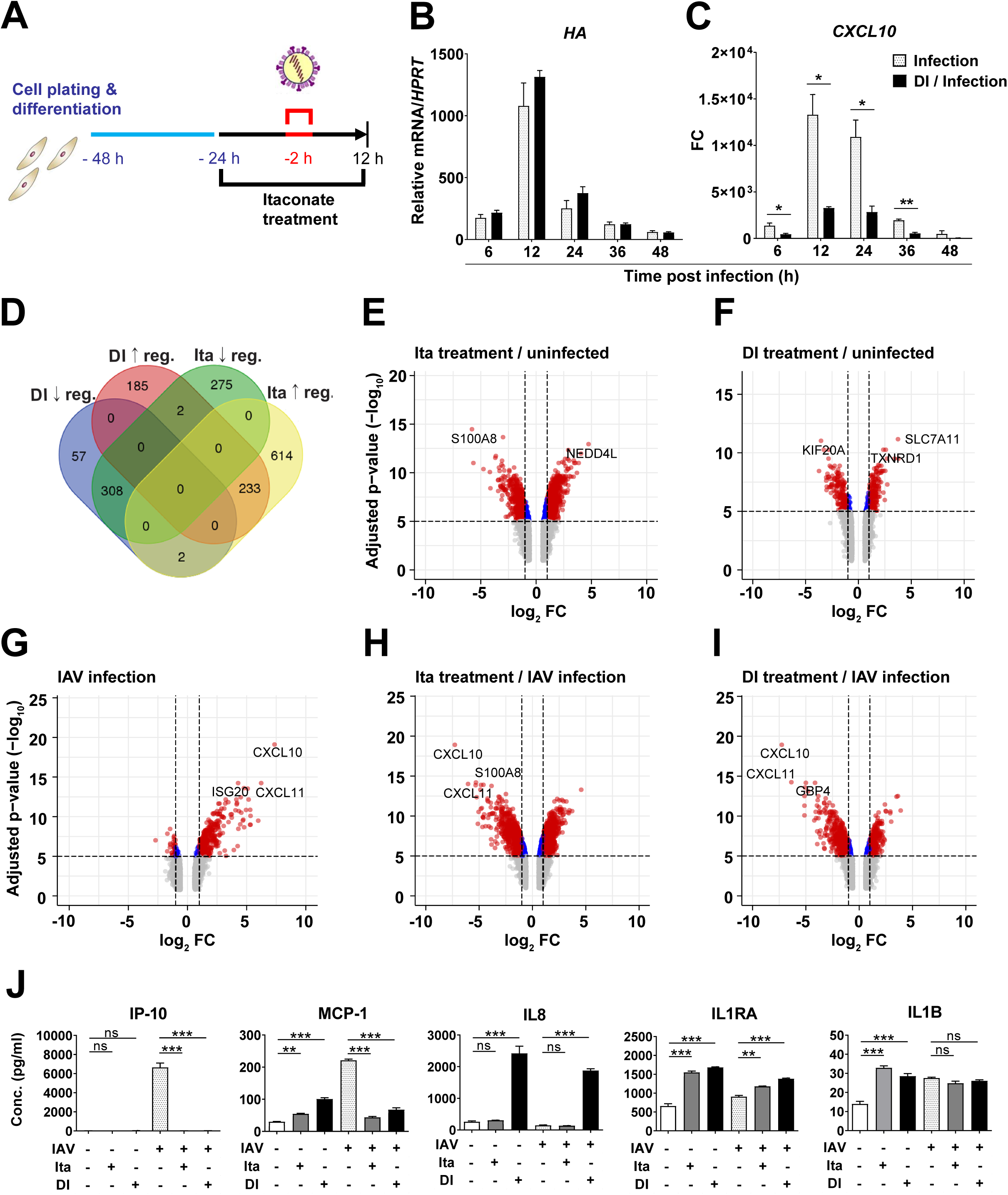
Effects of itaconate and DI on cellular inflammation due to IAV infection of dTHP-1 cells. **A.** Outline of experiment. dTHP-1 cells were treated with itaconate or DI overnight, or left untreated, and then infected with IAV. Dose-response curves of effects of a range of itaconate and DI and concentrations on *HA* and *CXCL10* mRNA expression are shown in Fig. S1**. B**-**C**. 48 h time course, DI = 0.25 mM (n=3). mRNA expression (RT-qPCR) of *HA* (**B**) and *CXCL10* mRNA (**C**) *p<0.05; **p<0.01; ***p<0.001 (unpaired t-test). **D-I.** Global transcriptomic changes (microarray analysis) due to itaconate (25 mM) and DI (1 mM) treatment of uninfected and IAV-infected dTHP-1 cells (n=3). **D.** Venn diagram showing unique and common differentially expressed genes by itaconate and DI treatment of infected cells. **E-I.** Volcano plots showing differential mRNA expression in uninfected dTHP-1 cells under itaconate and DI treatment (**E-F**) and infected dTHP-1 cells with or without itaconate or DI treatment (**G-I**). The 2-3 most significantly differentially expressed mRNAs are identified by labels. **J.** Effects of itaconate (25 mM) and DI (1 mM) treatment on levels of inflammation-related polypeptides in dTHP-1 cell supernatants 12 h after IAV infection or mock infection. Mean ± SEM (n=3). *p<0.05; **p<0.01; ***p<0.001 (One-way ANOVA, Tukey’s multiple comparison test).

Analysis of enriched GO terms revealed that IFN-I signaling was strongly induced in the infected cells, which was prevented by treatment with itaconate and DI (Fig. 4). This analysis also revealed that both compounds activated metabolic processes, whereas itaconate also modulated processes relating to differentiation and membrane signaling. A KEGG pathway analysis additionally revealed induction of classic pro-inflammatory pathways such as *TNF signaling*, *TLR receptor signaling*, and *chemokine signaling* by infection, all of which were depleted by itaconate and DI treatment (Fig. S5). Taken together, these results revealed strong anti-IFN and anti-inflammatory effects of itaconate and DI, but also suggested that they exert other important effects on cell metabolism and differentiation, some of which differ between the compounds.

**Figure 4.**
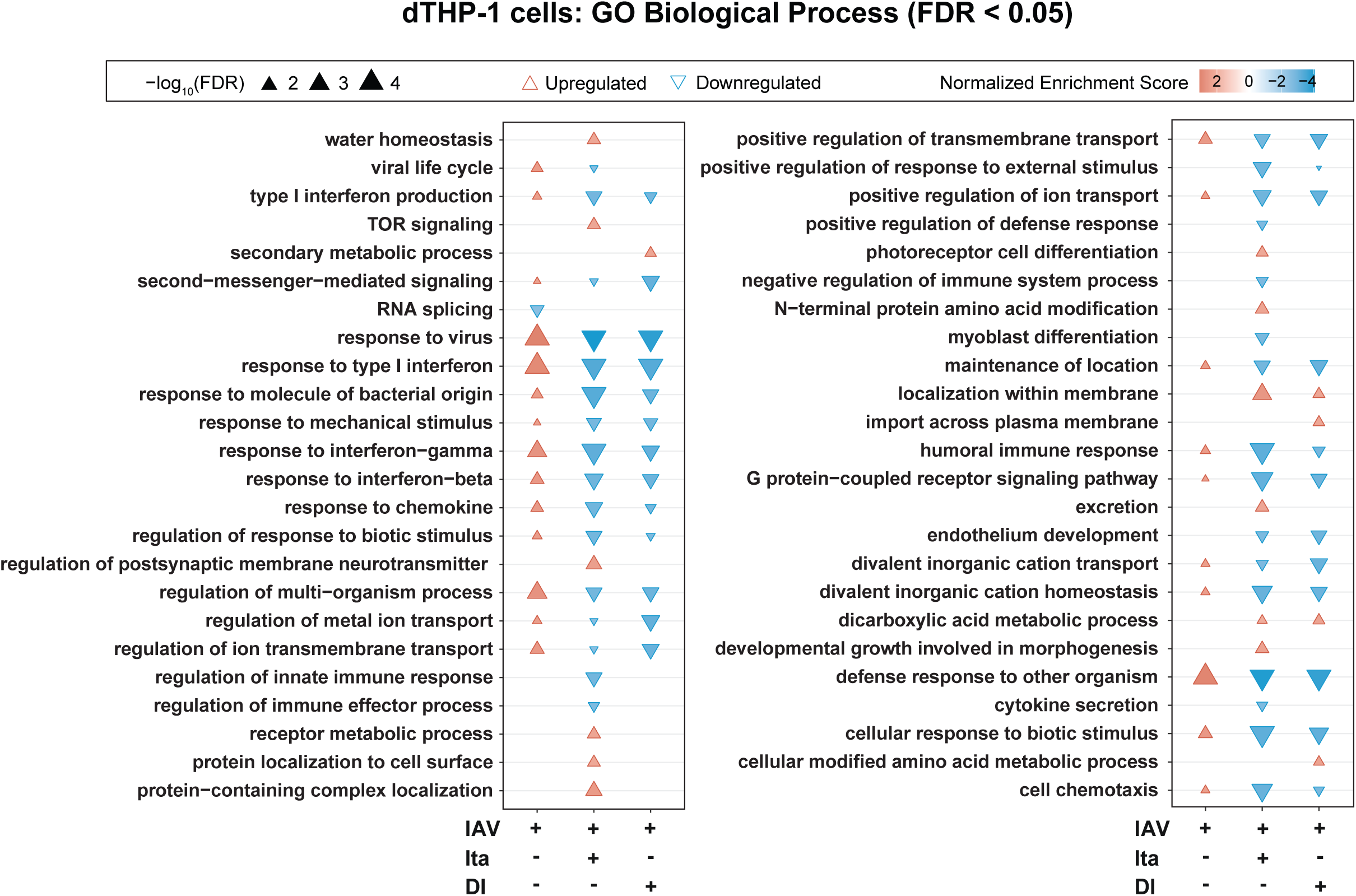
Itaconate and DI treatment leads to a global downregulation of IFN-I signaling in IAV-infected dTHP-1 cells. Enrichment analysis of gene ontology (GO Biological Process) terms based on the microarray data shown in Fig. 3, using DEGs (p<0.05, FC >|1.5|) as input. GO terms (FDR ≤0.05) reveal major induction of antiviral and inflammatory responses by infection, which are decreased by both itaconate and DI treatment.

### Anti-inflammatory effects of ectopically expressed itaconate, and exogenous itaconate and DI on IAV-infected respiratory epithelial cells

We then tested whether the effects of itaconate and DI could also be seen in cells not endogenously expressing *ACOD1*. For this purpose, we studied the respiratory epithelial cell line A549, which supports productive IAV infection and does not express *ACOD1*. As expected, infection of A549 cells did not result in induction of *ACOD1* mRNA, but transient transfection with an *ACOD1*-expressing plasmid led to significant *ACOD1* mRNA expression (Fig. 5A), which was accompanied by appearance of substantial concentrations of IA (Fig. 5B). While there was no effect on viral *HA* mRNA expression, transfection with the *ACOD1*-expressing vector led to a marked reduction in infection-associated *CXCL10* induction (Fig. 5C,D). A preliminary treatment experiment showed that (as opposed to THP-1 cells) DI was cytotoxic at 1.0 mM in A549 cells, and the 0.5 mM concentration was therefore used. Treatment with DI resulted in a modest reduction of *HA* expression early in infection and a marked reduction of *CXCL10*, *IL6*, and *IFNB* expression (Fig. 5E-H). Exogenous itaconate, too, reduced *CXCL10* mRNA, but less efficiently (Fig. S6). Taken together, these results obtained with A549 cells suggest that immunomodulatory effects of itaconate and DI are independent of the ability of the cell type to express endogenous *ACOD1*, and that similar effects on IFN-directed gene expression can be obtained with ectopic, endogenous synthesis of itaconate and exogenously added itaconate and DI.

**Figure 5.**
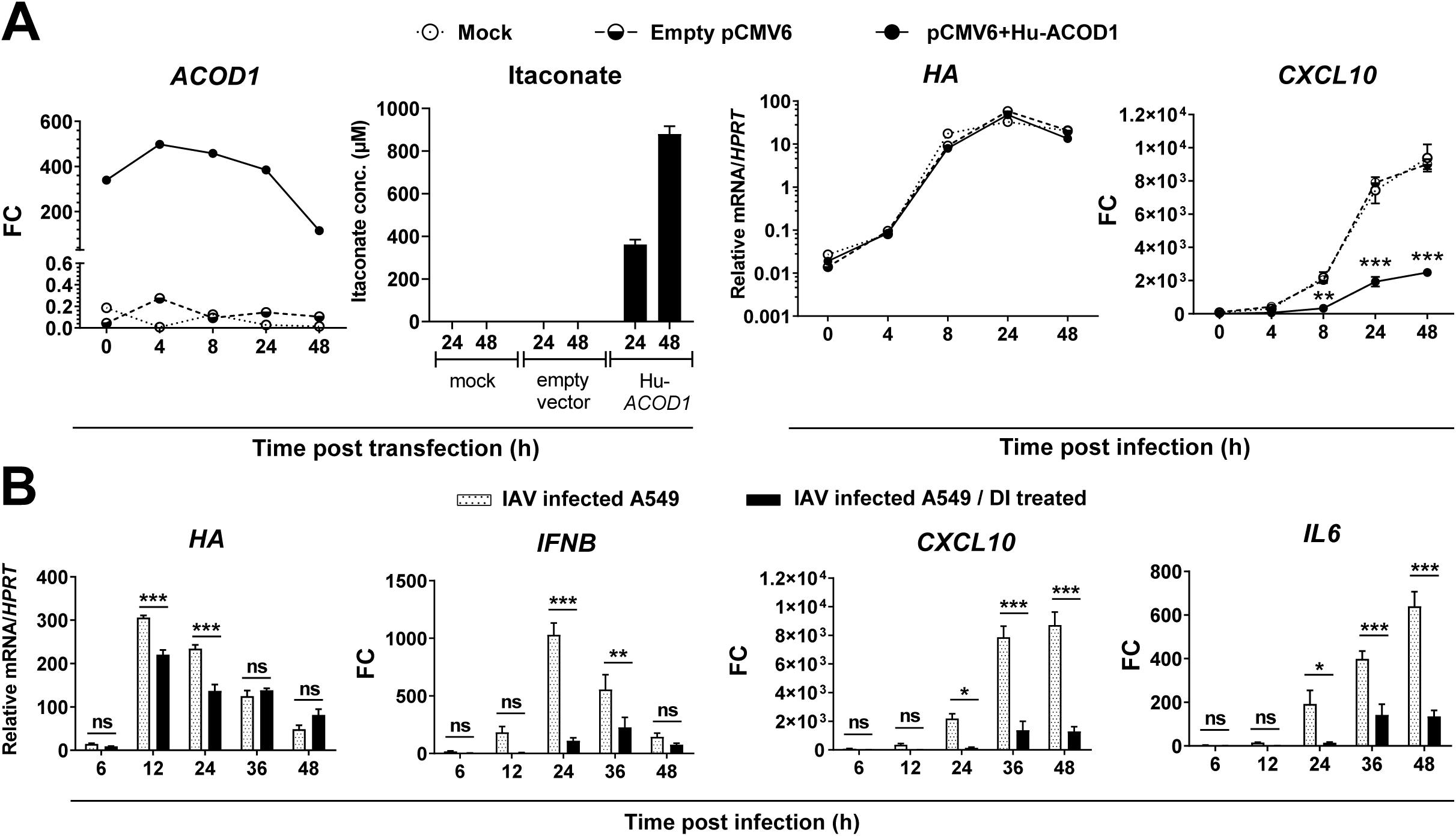
Effect of overexpression of ACOD1 and exogenous treatment with DI on IAV-infected A549 cells. **A.** A549 cells were transfected with pCMV6-Hu-ACOD1, pCMV6 empty vector, or mock transfected for 48 h and then infected with IAV for 48 h (n=3). Expression of the indicated mRNAs was measured by RT-qPCR, itaconate concentrations by LC-MS/MS. *p<0.05; **p<0.01; ***p<0.001 (2-way ANOVA, Tukey’s multiple comparisons test) **B.** A549 cells were treated with 0.25 mM DI overnight, infected with IAV for 48 h (n=3), and expression of the indicated mRNAs measured by RT-qPCR. Mean ± SEM (n = 3). *p<0.05; **p<0.01; ***p<0.001 (2-way ANOVA, Sidak’s multiple comparisons test).

Next, we assessed effects of itaconate and DI on cellular transcriptomes in IAV-infected A549 cells. PCA revealed the expected normalization of gene expression due to DI treatment, but also increasing effects of the compounds on overall cell homeostasis (Fig. S7). As in THP-1 cells, hierarchical clustering analysis revealed clear clustering of each of the four groups and the close relationship of the uninfected and DI-treated infected cells (Fig. S8). However, the anti-inflammatory effect of itaconate was weaker than that of DI, and at the higher (40 mM) concentration, the itaconate signature became so dominant that the itaconate-treated infected group now formed its own clade (Fig. S9). IAV infection enriched GO terms related to IFN-I and –II responses as well as other antiviral and pro-inflammatory terms, which could all be reduced to near baseline by treatment with DI or itaconate (Fig. 6). Visualization of differentially expressed IFN-related mRNA also showed that, in spite of its overall lower effect on pro-inflammatory signaling than DI, itaconate treatment at the lower (20 mM) concentration did have a considerable impact on IFN-I expression (Fig 6B). In addition, itaconate uniquely inhibited *NFKB signaling* and *fatty acid metabolism*. KEGG-pathway analysis additionally revealed depletion of other pro-inflammatory pathways, including *TNF signaling* and *necroptosis,* by the treatments (Fig. S10).

**Figure 6.**
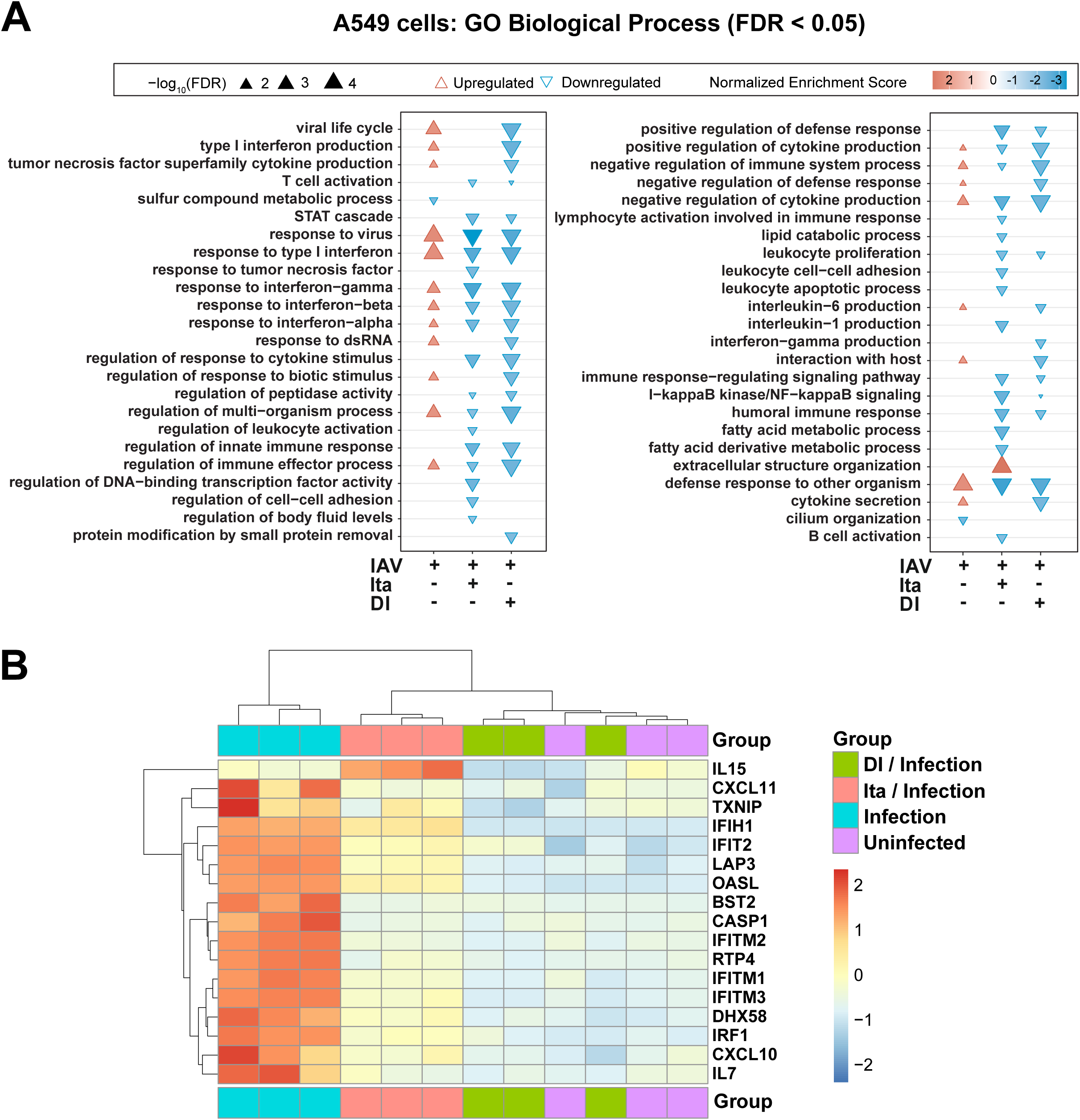
GO term enrichment analysis of itaconate and DI reveals global effects on IFN-signaling and cellular inflammation in A549. A549 cells were, in the case of treatment, incubated overnight with DI (0.5 mM) or itaconate (20 mM), and then infected with strain PR8M. Gene expression was determined by microarray analysis 24 h p.i. (n=4). **A**. GO enrichment (FDR <0.05) analysis using DEGs (p<0.05, FC >|1.5|) as input. **B**. Hierarchical cluster analysis illustrating downregulation of a subset of genes involved in IFNA/B signaling.

### Anti-inflammatory effect of itaconate and DI in a human lung tissue explant (HLTE) model of IAV infection

To test whether the *ACOD1*/itaconate axis and the anti-inflammatory effects of itaconate and DI are functional in the end-organ of IAV infection in humans, we then used a human lung tissue culture model. Indeed, infection with IAV led to brisk transcription of viral *HA* and host *CXCL10* mRNA in a 72 h time course (Fig. S11), whereas *ACOD1* mRNA levels did not change significantly. Adding itaconate and DI reduced expression of *CXCL10* mRNA in tissue and IP10 protein in culture supernatant, and itaconate additionally reduced ISG15 mRNA in tissue, whereas viral titers in the supernatant were unaffected (Fig. 7). Re-analysis of a published dataset from a model using a different IAV strain (H3N2) and histologically normal lung tissue (Matos, Wunderlich et al., 2019) revealed a pronounced induction of *ACOD1* and *TNFAIP3* expression, but (as opposed to *in vivo* infection of mouse lung) not induction of *HMOX1* or *2* (Fig. S12).

**Fig. 7.**
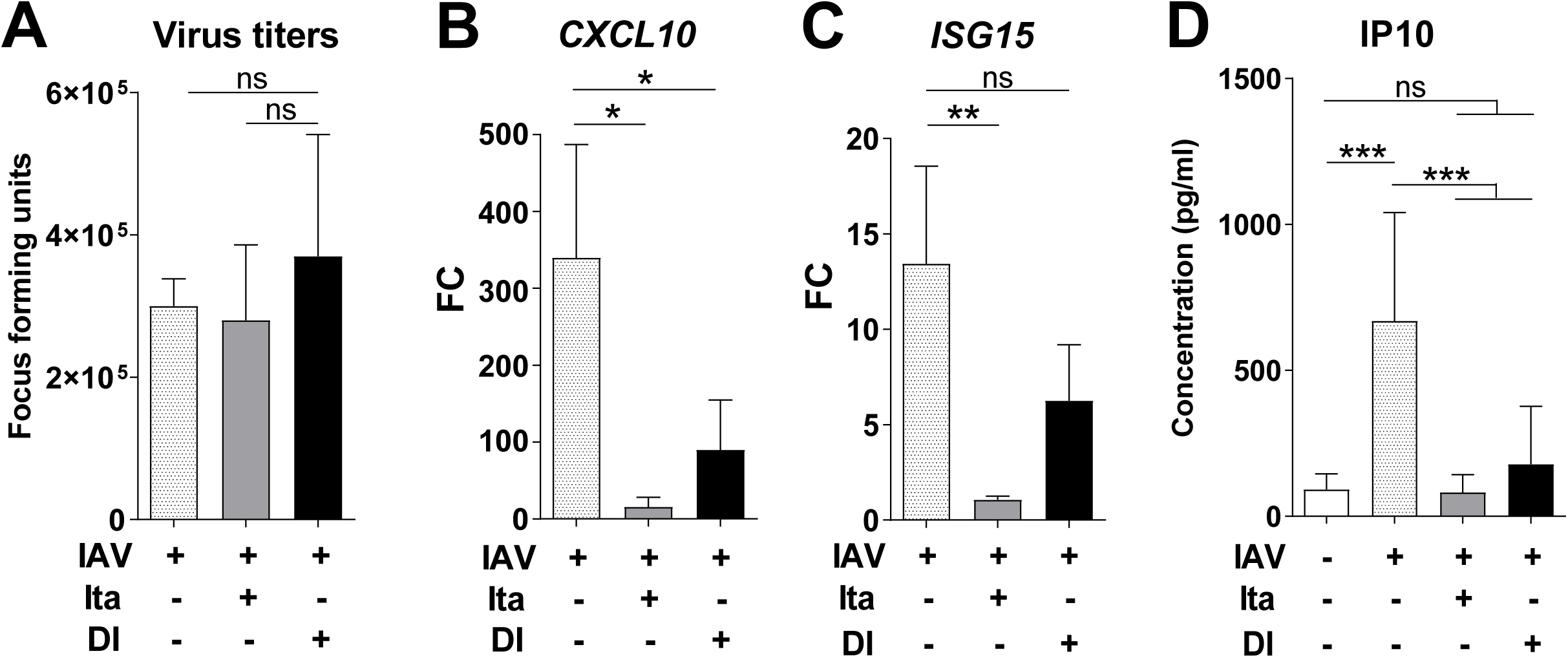
Anti-inflammatory effect of itaconate and DI on IAV-infected human lung tissue explants. Human primary lung tissue from patients with emphysema and pulmonary arterial hypertension was incubated overnight with itaconate (25 mM), DI (1 mM), or medium only and was then infected with IAV for 24 h (n=5 donors, 3 tissue pieces per donor). **A**. Viral titers in supernatant (focus forming assay). **B-D**. *CXCL10* mRNA (**B**), and ISG15 (**C**) mRNA in tissue (RT-qPCR); IP10 concentration in supernatant (EIA) (**D**). Mean ± SEM (n=6 donors, 3 tissue pieces per replicate). *p<0.05; **p<0.01; ***p<0.001 (Mann-Whitney U test).

### Anti-inflammatory effects of itaconate, DI, and 4OI on IAV-infected PBMC

PBMC play important roles in influenza infection as they contain cells of innate and adaptive immunity, and because circulating monocytes (which can sustain a nonproductive influenza virus infection) are considered an important source of cytokines that support systemic spread of inflammation (Duan, Hibbs et al., 2016). We therefore investigated effects of the compounds on IAV infected PBMCs both at the bulk and single cell levels. Analysis with targeted PCR- and immuno-assays demonstrated vigorous transcription of viral RNA, induction of *ACOD1* mRNA, and a brisk pro-inflammatory response both at the mRNA and protein level (Fig. 8A,B). The induction of *ACOD1* mRNA expression in peripheral blood in IAV infection was also corroborated by reanalysis of a published dataset from whole blood from patients with moderate and severe influenza (Fig. S13). The effects of itaconate were mixed in that there was no significant change in *ACOD1*, *IFNB1*, and *TNF* mRNA expression in cells or of IL1B protein in supernatants. In contrast, DI and 4OI led to a marked reduction of IFN and pro-inflammatory cytokine expression at the mRNA and protein level, as well as of *ACOD1* mRNA. As exemplified by the effect of DI and 4OI on *TNF* and *ACOD1* expression, the treatments reduced expression of some mRNA targets even in uninfected cells, suggesting that they also reduced some baseline inflammation. Of note, 4OI in addition led to a pronounced (95%) reduction in *HA* transcription (p=<0.0001), indicating inhibition of viral RNA replication. PCA of bulk PBMC transcriptomes revealed strong effects of the compounds on cell transcriptomes in general, which led to one outlying DI-treated sample and two itaconate-treated samples, whereas all four 4OI-treated samples formed a clearly discernible group (Fig. S14). The most significant transcriptome changes were driven by 4OI, which had major effects on many genes that were not regulated by IAV infection or itaconate or DI treatments (Fig. S15). Nonetheless, induction of *HMOX1* by DI in IAV-infected samples was evident. Functional enrichment analysis of GO terms based on the transcriptomes revealed that infection induced the expected antiviral responses, including *response to IFN-I*, as well as several other inflammation-related terms (Fig. 8C). At least one of the three compounds prevented activation of each of the infection-associated terms. However, consistent with its lack of effect on *IFNB* expression (Fig. 8), itaconate did not affect *response to IFN-I* or *influenza A*. Several terms that were depleted by the compounds in infected PBMC were not enriched by infection alone, suggesting that the compounds diminished also a baseline activation of the PBMC. Interestingly, 4OI treatment strongly enriched terms relating to chromatin conformation, which turned out to be driven by upregulation of histone deacetylase epression. A KEGG pathway analysis essentially confirmed the findings of the GO term analysis (Fig. S16), including that 4OI enriched pathways relating to chromatin structure such as *Alcoholism*. Taken together, these results consolidated our above findings that itaconate and derivatives can modulate inflammation due to IAV infection, but also suggested that DI and 4OI may have more robust anti-IFN and anti-inflammatory effects than itaconate. Considering the variable anti-inflammatory effects of itaconate and the miscellaneous effects of 4OI on gene expression in this model, we subsequently focused on DI to decipher effects at the single-cell level.

**Fig. 8.**
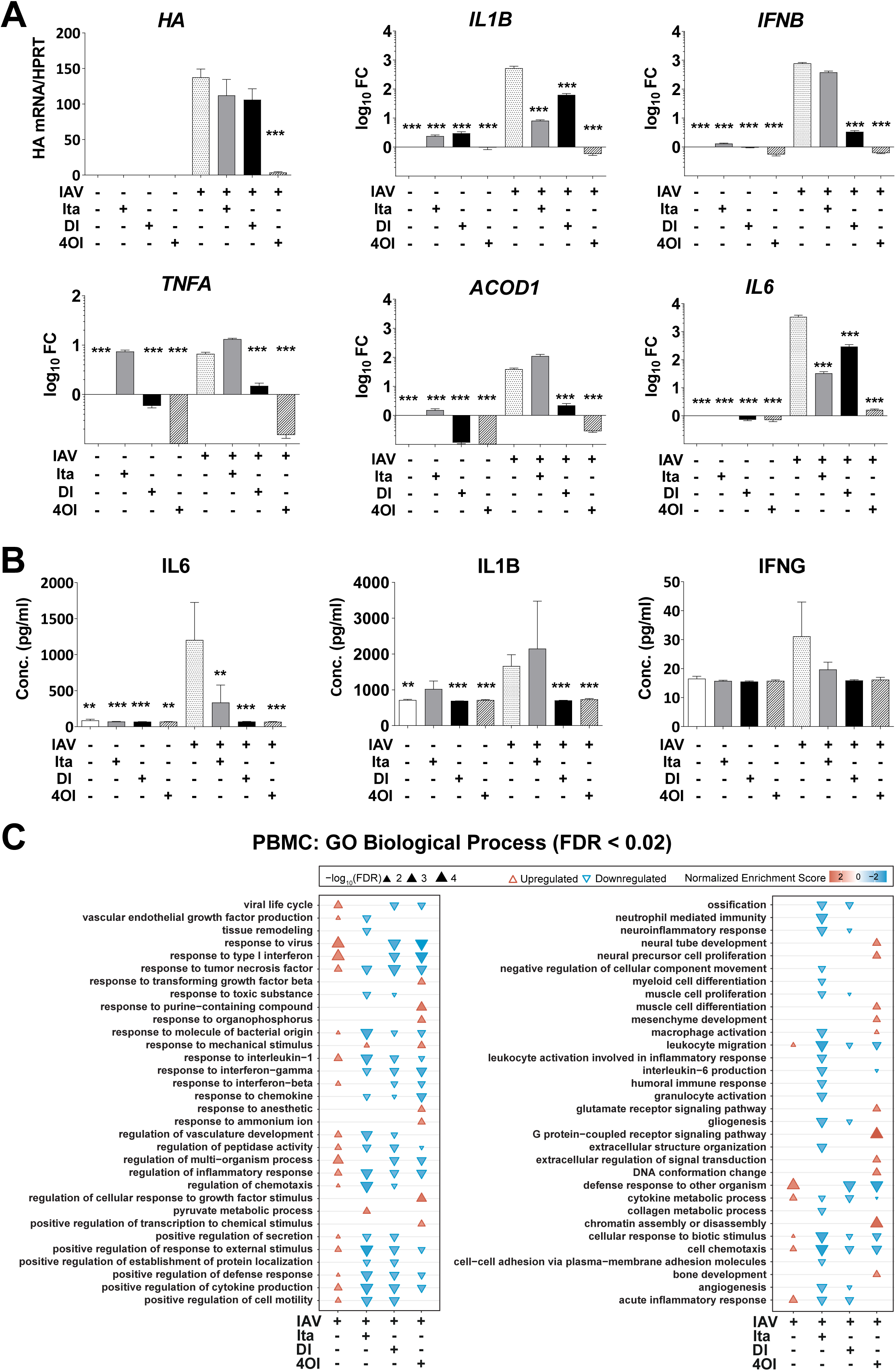
Anti-inflammatory effects of itaconate, DI, and 4OI on IAV-infected human PBMCs. PBMCs were isolated from freshly donated human blood and then infected with IAV for 12 h in the presence or absence of itaconate (10 mM), DI (0.5 mM), or 4OI (25 µM) without preincubation (7 donors, 3 replicates). **A.** Expression of the indicated mRNAs in PBMC (RT-qPCR). **B.** Concentrations of the indicated proteins in PBMC supernatants (ELISA). Mean +/− SEM (6 donors, 3 replicates per donor). *p<0.05; ***p<0.01; ***p<0.001 (1-way ANOVA). **C.** GO enrichment analysis of effects of IA, DI, and 4OI on IAV-infected PBMC. Microarray analysis of PBMC RNA from four of the donors featured in **A-B**. RNA was pooled from the three replicates of each donor, resulting in 4 samples per group. GO term analysis was performed on DEGs (p<0.05, FC>|1.5|) and terms with an FDR <0.02 in at least one condition are shown.

### Single-cell RNAseq of PBMC identifies monocytes as the main PBMC host cell of IAV infection, sole source of *ACOD1* expression, and predominant target of immunosuppressive effects

Thirteen cell types could be identified in the PBMC scRNAseq data set, but we focused the analyses on the overarching five cell types: CD4+ and CD8+ T cells, B cells, NK cells, and monocytes (Fig. 9A,B). Monocytes constituted the least numerous cell type, and DI treatment of control and infected cells led to a further reduction (Fig. S17). It was not possible to discern whether this was due to decreased survival or a technical artefact. Monocytes were essentially the only cell type expressing viral RNA, as only negligible amounts of viral transcripts were detected in lymphocytes and NK cells. DI treatment appeared to result in a slight increase in viral RNA expression in monocytes (Fig. 9C). IAV infection upregulated expression of *ACOD1* exclusively, and *CXCL10* predominantly, in monocytes, whereas *TNFAIP3*, *IFIT1,* and *ISG15* were upregulated also in the NK, T and B cell compartments (Fig. 9D). DI treatment markedly reduced expression of *ACOD1*, *IFIT1,* and *CXCL10*, but –interestingly– it reduced *TNFAIP3* and *ISG15* (albeit weakly) expression only in monocytes. Consistent with the observation that they constituted the main PBMC host cell, IAV infection triggered the most vigorous transcriptomic host response in monocytes, which was driven by the expected IFN responses (Fig. 9E). However, discernable differential expression (mostly of IFN-related RNAs) was also observed in the other cell types. CD8+ T cells are shown as a representative example in Fig. 9F, the other cell types in Fig. S18. Monocytes were the only cell type that mounted a transcriptomic response to DI treatment of uninfected cells. In infection, treatment with DI led to a global downregulation of IFN-related transcripts in monocytes and also downregulation of a subpopulation of IFN-related transcripts in lymphocytes and NK cells. When considering all DEGs irrespective of magnitude of differential expression, DI exerted comparable effects on monocytes and T cells, and only somewhat less on B cells. However, when considering only transcripts with FC >|2|, a preferential effect on monocytes became apparent (Fig. 9G,H).

**Fig. 9.**
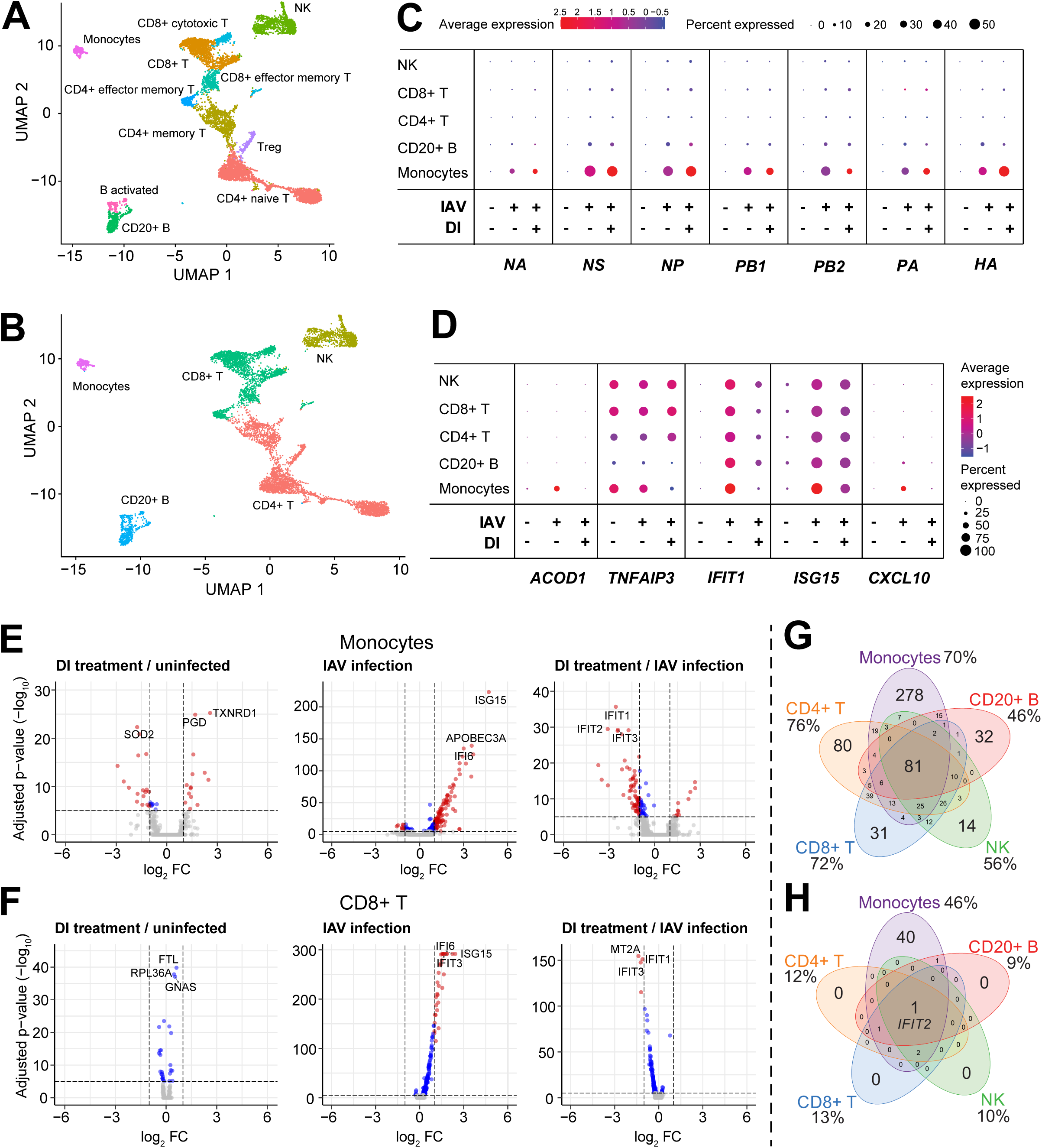
Single-cell transcriptomic responses of PBMC to IAV infection and DI treatment. PBMCs were isolated from freshly donated human blood (one donor), and then infected with IAV for 12 h with or without DI (0.5 mM) in the medium. **A-B**. UMAP plots identifying 13 cell types (**A**) that make up the five major cell types (**B**) studied. The RNA cell markers used are listed in Table 3. **C-D.** Differential mRNA expression in uninfected, infected and infected/DI-treated PBMCs. **E.** Differential mRNA expression in monocytes in the indicated paired comparisons. Major transcriptional reprogramming due to IAV infection is evident, which is largely prevented by treatment with DI. **F.** Differential mRNA expression in CD8+ T cells in the indicated paired comparisons. Reprogramming in the presence of IAV is much less, but a DI treatment effect is evident. **G-H.** Venn diagrams based on RNAs differentially regulated (FDR <0.05) by both viral infection and DI treatment in each of the 5 major cell type, either irrespective of FC (**G**) or FC >|2| (**H**).

**Table 1.** Scoring system to assess inflammation in IAV-infected mouse lung.

|  |  |  |  |  |
| --- | --- | --- | --- | --- |
| **Parameter** | 0 | 1 | 2 | 3 |
| Severity of infiltration of inflammatory cells | Not present | Mild | Moderate | Severe |
| Area involved | No changes | 1-30% | 30-70% | >70% |

**Table 2.**
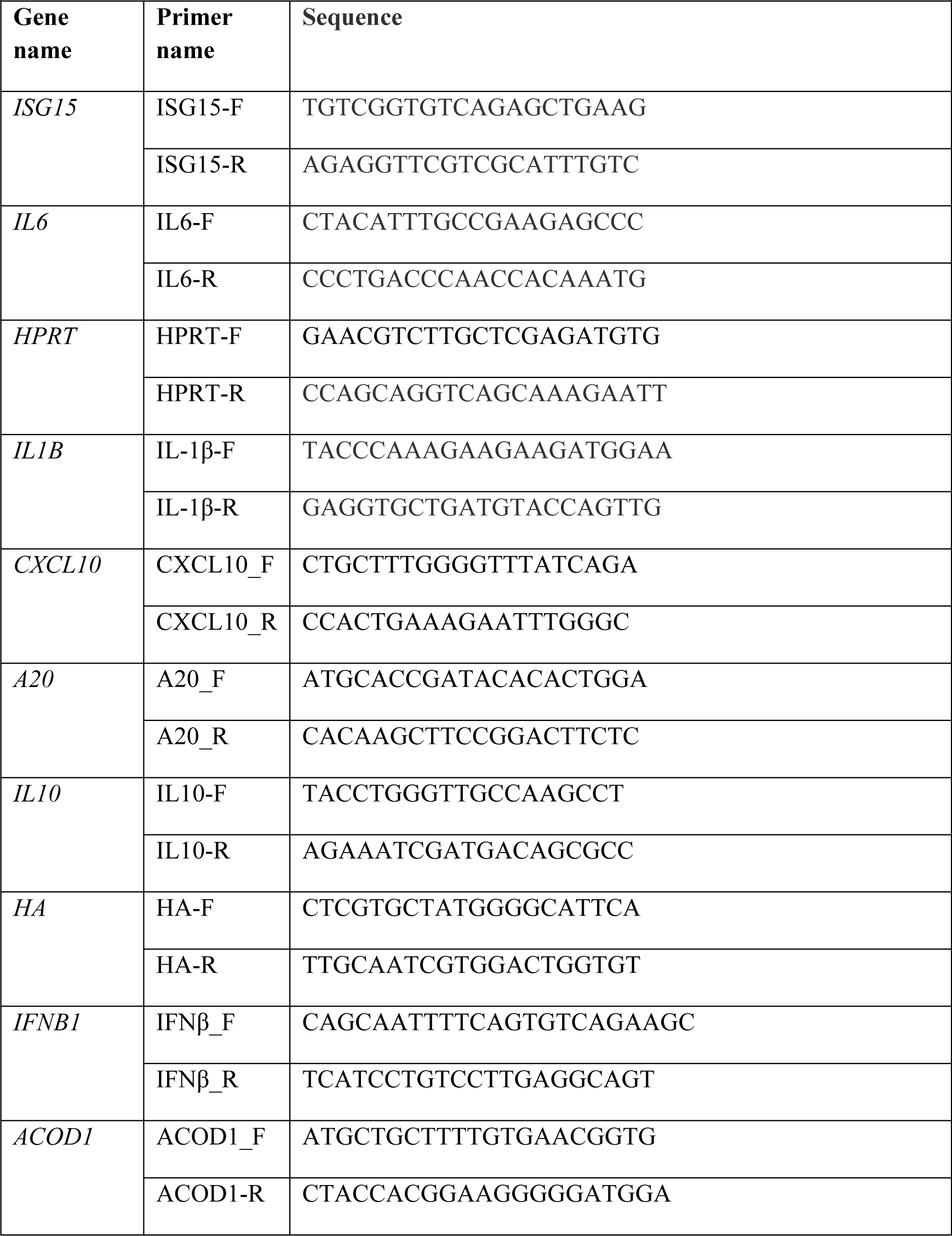
List of RT-qPCR primers.

**Table 3.** mRNA markers used to define PBMC cell subpopulations in scRNAseq.

|  |  |  |  |  |  |
| --- | --- | --- | --- | --- | --- |
| **Cell type** | **Sub-type** | **Marker(s)** | **Sub-sub type** | **Marker(s)** | **Ref.** |
| T cells | CD4 | *CD3D, IL7R* | CD4 naïve | *LEF1, ATM, SELL, KLF2, ITGA6* | (Butler, Hoffman et al., 2018, Cano-Gamez, Soskic et al., 2020, Szabo, Levitin et al., 2019) |
|  | CD4 |  | Effector memory | *KLRB1 IL7R* | (Butler et al., 2018, Cano-Gamez et al., 2020) |
|  | CD4 |  | CD4 memory | *PASK* | (Butler et al., 2018, Cano-Gamez et al., 2020) |
|  | CD4 |  | Treg | *FOXP3* | (Li, Li et al., 2015) |
|  | CD8 | *CD3D, CD8A* | Effector memory | *GZMK* | (Butler et al., 2018, Chu, Berner et al., 2020) |
|  | CD8 |  | Cytotoxic | *GNLY* | (Butler et al., 2018, Szabo et al., 2019) |
| B cells |  | *MS4A1* | Activated B cells | *MIR155HG* | (Butler et al., 2018) |
| NK cells |  | *NKG7*<br>*GNLY* |  |  | (Butler et al., 2018) |
| Monocytes |  | *CD14, LYZ* |  |  | (Butler et al., 2018) |

GO term enrichment analysis of the monocyte scRNAseq data confirmed the major enrichment of IFN-related processes by infection and their depletion by DI treatment, which had been seen in all models in this study. In addition, infection broadly dampened RNA metabolism, ribosome assembly, and protein synthesis and export, which was consistent with the cytostatic effects of type I IFN. DI treatment reversed all these effects and also stimulated some in uninfected cells (Fig. 10A). A KEGG enrichment analysis of these monocyte data confirmed the depletion of IFN-related pathways (*influenza, measles, cytosolic DNA sensing pathway*) by DI treatment, but also of other pro-inflammatory pathways (*TLR signaling, cytokine-cytokine receptor interaction, NF-kB signaling*) in infected and uninfected cells (Fig. S19). Of note, DI treatment apparently stimulated ribosome function in both infected and control monocytes.

**Figure 10.**
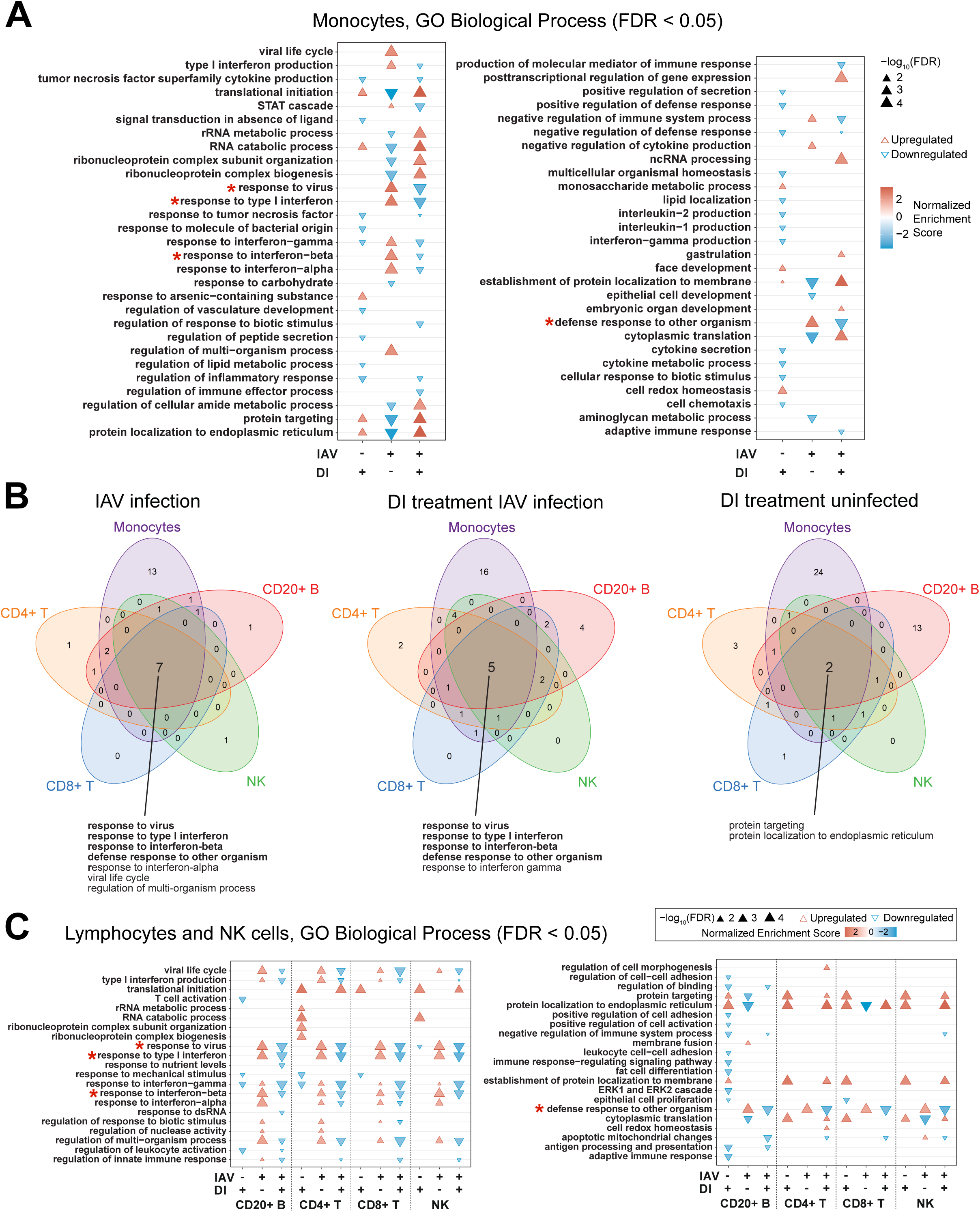
GO enrichment analysis at the single cell level of DI effects on uninfected and IAV-infected PBMC. Analysis based on the scRNAseq data shown in Fig. 9. Go terms that are enriched/depleted in all cell types by infection and DI treatment are marked with a red asterix in A and C. **A.** GO enrichment analysis of monocytes. **B.** Venn diagrams illustrating GO terms that are commonly or uniquely regulated (FDR <0.05) in monocytes, CD4 T cells, CD8 T cells, B cells, and NK cells; due to infection alone, DI treatment of infected cells, or DI treatment of uninfected cells. Go terms that are enriched/depleted in all cell types by infection and DI treatment are printed in bold. **C**. Go term analysis of CD4 T cells, CD8 T cells, B cells, and NK cells.

The Venn diagrams in Fig. 10B illustrate that monocytes are the cell type in which the most biological processes were activated by infection and depleted by DI treatment, but reduction of antiviral and pro-inflammatory responses was a common theme in the other cell types as well (Fig. 10C). Indeed, GO term analysis identified four major IFN-related processes as a central functional network that was upregulated by infection and downregulated by DI treatment during infection in all five cell types (Fig. 10B). A marked stimulation of protein synthesis and export by DI was also apparent in the uninfected lymphocytes and NK cells. Finally, in order to identify a common transcriptomic network we performed GO and KEGG pathway enrichment analyses of the 81 transcripts that were commonly differentially expressed in IAV infection and regulated in the opposite direction by DI treatment in all five cell types (center of the Venn diagram in Fig. 9G). While the GO analysis by and large confirmed the findings shown in Fig. 10B, the KEGG pathway analysis revealed a major effect of DI treatment on *allograft rejection*, *graft-vs.-host disease*, and *antigen processing and presentation,* suggesting a broader interference with immune cell activation by DI (Fig. S20).

## Discussion

In this first dedicated study on the impact of itaconate and derivatives on host responses to IAV infection, we found that (i) the endogenous ACOD1/itaconate axis limits pulmonary inflammation due to the infection in a murine model, (ii) ectopically expressed ACOD1 possesses anti-IFN activity in an epithelial ell line that normally does not express this enzyme, and (iii) that exogenously applied itaconate and/or derivatives restrict IFN-responses and modulate classic pro-inflammatory cytokines across all models tested.

### Anti-inflammatory effects of itaconate and derivatives

A major suppression of IFN and some pro-inflammatory cytokine responses was the recurrent theme in all models, i.e. immortalized cells, primary cells, and human lung tissue. This is fully consistent with the early reports that DI has strong anti-inflammatory properties (Lampropoulou et al., 2016), which was subsequently corroborated in diverse models of inflammation and infection, also using 4OI (e.g., (Gu et al., 2020, Xu, Jiang et al., 2020a, Zhao et al., 2019)). We used itaconate and DI in parallel in several models, which enabled comparing their relative effectiveness. Overall, results with DI were most consistent, and in spite of its higher toxicity on A549 cells than on dTHP-1 cells, the lower dose that had to be used on A549 cells still possessed strong anti-inflammatory effects. Itaconate effects were less consistent and much higher doses had to be used to achieve anti-IFN effects similar to DI. Itaconate did not reduce expression of *IFNB1*, *TNF* mRNA or IL1B protein in PBMC. The lack of *TFN* repression by itaconate in that model is consistent with previous reports that it usually does not affect TNF-α level (Bambouskova et al., 2018). Recent work has also revealed the potential of itaconate to actually enhance IL1-β release (Muri, Wolleb et al., 2020). Of note, at higher doses, itaconate and strong electrophilic compounds can actually increase inflammation due to CASP-8 dependent inflammasome activation, and this effect is magnified when the cells are not pre-incubated with the compound before adding the pro-inflammatory stimulus (Muri et al., 2020), as we did in the PBMC infections. However, these authors did not investigate effects on IFN suppression. 4OI demonstrated the expected potent anti-inflammatory effects, but also broad effects on processes relating to, e.g., chromatin structure, the potential effects of which on cell homeostasis and cell-pathogen interactions will require further study.

### Anti-inflammatory effect of ectopic synthesis of itaconate in A549 cells

Despite the multitude of studies on effects of exogenous administration of itaconate derivatives on cellular inflammation, it had not been tested whether ectopic synthesis of itaconate in a cell type that does not naturally express *ACOD1* has similar effects. Our transfection study (Fig. 5) demonstrated that ectopic, endogenously synthesized itaconate exerts very similar anti-IFN effects as exogenously added itaconate or DI. Of note, the achieved itaconate concentration is very close to that measured in LPS/IFNG stimulated dTHP-1 cells (Chen et al, unpublished data), suggesting that it is physiological.

### Cell-type specific and -independent anti-inflammatory effects of itaconate and DI

A particular strength of our study lies in the use of scRNAseq to identify major target cells in peripheral blood. Firstly, this analysis identified monocytes as the major PBMC type that is infected by IAV. Secondly, we found that on one hand DI can attenuate IFN-responses in lymphocytes and NK cell as well, but that the immunosuppressive effect is greatest on monocytes. Taken together, these results suggest that even though pharmacological use of itaconate and derivatives would modulate host responses in a diverse array of cells, monocytes/macrophages likely constitute a major functional hub in the genesis of inflammation in IAV infection and subsequent treatment responses to itaconate-based adjunct treatments.

### Antiviral effect of 4OI

Olagnier et al. recently reported that Nrf2 activators including 4OI and dimethyl fumarate limit inflammation in cellular models of SARS-CoV-2 infection (Olagnier, Farahani et al., 2020) in an NRF2-dependent manner. Of note, as opposed to influenza viruses, infection with SARS-CoV-2 suppresses the NRF2 pathway, which can be reversed in part by treatment with Nrf2 activators. These authors in addition observed an antiviral effect of 4OI, which proved to be independent of NRF2- and type I IFN-signaling. Our finding of a remarkable suppression of IAV RNA replication in PBMC corroborates the notion that 4OI has strong antiviral effects in addition to its well-documented anti-inflammatory properties (Olagnier et al., 2020). Further research is needed to validate our finding in other cell types, as we tested it in PBMC only. Nonetheless, the gene expression analysis of itaconate, DI- and 4OI-infected PBMC identified several processes/pathways uniquely regulated by 4OI. Further research should focus on assessing whether any of them could contribute to the antiviral effect of 4OI.

### Infection-independent effects on target cells

By using relatively high doses of itaconate and DI and assessing effects of the compounds on uninfected cells or their effects on processes that were not affected by infection alone, our study also provides a first comprehensive look at global effects of the compounds on target cells. In particular, itaconate and 4OI exerted pronounced, broad effects on gene expression in A549 cells and PBMCs, respectively. Clearly, it will be of great interest to investigate how these effects relate to differences in toxicities on different cell types, or to differences in their effects on virus-host interactions.

### Importance of endogenous ACOD1 expression

In the mouse model, we observed a more severe clinical phenotype and higher inflammation in *Acod1^-/-^* mice. This agrees with work in a mouse model of *M. tuberculosis* infection, where inflammation and mortality were higher in the *Acod1^-/-^* animals (Nair et al., 2018). However, IAV infection was also assessed in that study, and deleting the *Acod1* gene did not affect weight loss or survival. Those authors used equal numbers of male and female mice, whereas we used female mice only. Further research is necessary into possible sex-specific effects of the *Acod1* gene in influenza infection. Of note, we found upregulation of *ACOD1* by IAV infection and its prevention by DI treatment in myeloid cells, i.e. dTHP-1 and peripheral blood monocytes, and upregulation of *ACOD1* mRNA in blood from patients with moderate and severe influenza infection as compared with healthy donors. Thus, induction of *ACOD1* expression (and thereby itaconate synthesis) likely is a common feature in human influenza. We have recently identified the active center of ACOD1 enzyme and found that naturally occurring loss-of-function mutations are extremely rare, suggesting that physiologic itaconate responses have been important in human evolution (Chen et al., 2019). It is therefore tempting to speculate that one beneficial role of an active ACOD1/itaconate axis is to reduce the risk of death or organ damage from infection-associated inflammation and that loss-of-function genotypes have been selected against during times of high infectious disease burdens.

### Clinical implications

The current COVID-19 pandemic has, once again, focused the research limelight on the deleterious effects of overshooting innate immune responses in acute viral infections. In a current report, Lee et al. found that intense IFN responses in PBMC from patients are a hallmark of both severe COVID-19 and influenza (Lee, Park et al., 2020). Of note, a recent systematic review of 45 studies on corticosteroid use for COVID-19 found a significant beneficial effect (van Paassen, Vos et al., 2020), whereas to date there are no clinically effective antivirals against SARS-CoV-2. Therefore, the concept of adjunct treatments to control deleterious host immune responses in viral infections is more pertinent than ever. Considering (i) the well documented cytoprotective properties of itaconate and derivatives and (ii) our findings of remarkable anti-inflammatory and anti-IFN effects in the absence of increased viral replication, itaconate derivatives would constitute a highly promising class of compounds for such applications. Considering the more reliable results obtained with DI as compared to unmodified itaconate, the lower required concentrations of DI and 4OI, but also the striking anti-viral effects of 4OI, it appears that chemical variants of itaconate, rather than its native form, will take the lead in further translational development to the bedside.

## Acknowledgements

We thank V. Kaever, H. Bähre and staff of the RCU Metabolomics at Hannover Medical School for support with the itaconate measurements, Elena Reinhard for assistance with the animal experiments, the HZI Animal Facility for expert animal care, the HZI Genome Analytics Platform for microarray and RNA sequencing, L. Gröbe for cell sorting, D. Jonigk and P. Braubach (Institute of Pathology, Hannover Medical School) for providing human lung tissue. We thank Hans Jörg Hauser (HZI) for helpful comments on the manuscript.

## Authors’ contributions

AS, AAI, CF, FC,NS, KS, MP, MT, and TE performed experiments and analyzed data, MW performed statistical analyses and prepared figures, RG performed the gene expression studies, MCP performed histological analyses, CAG, AM, SP and AM provided reagents and concepts to study design. All authors edited the manuscript and consented to its publication. FP oversaw the study and wrote the final draft of the manuscript.

## Data Availability

The gene expression data are available through GEO at accession number GSE162210, GSE162260, GSE162261 and GSE164922. These data are currently private and access can be given to reviewers. The data previously published in (Wilk et al., 2015) are available at GSE66040.

## Methods

### Cell culture

Cell lines were obtained from German Collection of Microorganisms and Cell Cultures GmbH (DSMZ), Braunschweig, Germany. The human myelomonocytic leukemia cell line THP-1 was propagated in RPMI 1640 medium (GIBCO^®^ Life Technologies™) supplemented with 10% fetal calf serum (FCS) and 2 mM L-glutamine. Cells (2.5×10^5^ cells/ml) were differentiated with 200 nM phorbol-12-myristate-13-acetate (PMA, Sigma-Aldrich, product no.P-8139) for 48 h, followed by incubation in fresh RPMI medium for another 24 hours (h). Human adenocarcinoma cells resembling type II alveolar epithelial cells (A549) were propagated in DMEM medium supplemented with 10% FCS and 2 mM L-glutamine.

### Primary cell isolation and differentiation

Primary human monocytes were isolated from buffy coats from healthy blood donors by density gradient centrifugation (Histopaque™, Sigma-Aldrich). For studies of monocytes and M1/M2-type macrophages, CD14+ cells were isolated by magnetic activated cell sorting (CD14+ Cell Isolation Kit; Miltenyi Biotec). 2×10⁵ monocytes were then differentiated into M1-type macrophages cells by incubation with 1000 U/ml GM-CSF (granulocyte macrophage-colony stimulating factor, CellGenix) and into M2-type macrophages by incubation with 100 ng/ml M-CSF (macrophage-colony stimulating factor, Miltenyi Biotec) in serum-free DC medium (CellGenix) for 5 days.

### Virus strains

A common reference strain of influenza A virus (PR8M, A/PuertoRico/8/34 H1N1, kindly provided by Stefan Ludwig (University of Münster, Germany) (Blazejewska, Koscinski et al., 2011) was used throughout. In the experiment shown in Fig. 2, we additionally used the clinical isolate Gi-WT (A/Giessen/6/2009 H1N1-WT) and the reassortant Gi-NS-PR8 containing the NS segment of the PR8 strain on the backbone of SOIV-WT, which replicates faster than the WT (Petersen et al., 2018b). Viruses were propagated in the chorio-allantoic cavity of 10-day-old embryonated eggs for 48 h at 37°C. Fluid from the chorio-allantoic cavity was collected and the virus was titrated by focus-forming unit (FFU) assay and stored in aliquots at - 80°C until use.

### Viral infections and treatment with itaconate, DI, and 4OI

2.5 × 10^5^ dTHP-1 or A549 cells were infected with the indicated IAV strain at an MOI of 1, cells were centrifuged at 300 g for 15 min and incubated at 37°C for 2 h. Cells were subsequently washed twice with PBS and fresh post-infection medium was added to the cells. Cells were re-incubated at 37°C for the indicated lengths of time. The respective amount of IA, DI, or 4OI was added to RPMI/DMEM complete medium and pH was adjusted to 7.5 with 1 M KOH solution. 4OI (100 mM) was dissolved in DMSO, and the final concentration of 4OI (25 µM) thus resulted in a 0.025% concentration of DMSO in the media. Media were filtered by using vacuum filters with 0.22 μm pore size (Millipore). Cells were incubated overnight with the compounds, and virus was then added for 2 h. Unbound virus was then removed by replacing the medium with fresh medium containing the compounds at the same concentrations. For PBMC infections, cells were isolated from 7 donors and seeded as 4×10^6^ cells/well in a 6 well plate and simultaneously infected for 12 h with strain PR8M (MOI=1) with or without IA (10 mM), DI (0.5 mM), or 4OI (25 µM). Thus, PBMC were not preincubated with the compounds before infection. Strain A/California/7/2009 (reagent 15/252, obtained from National Institute for Biological Standards and Control, London, UK) was used for the mouse infections, applying 5×10^5^ FFU (20 μL) by the intranasal route.

### Mouse model

All animal procedures were performed following guidelines from the Federation of European Laboratory Animal Science Associations. The study was approved by the regulatory authority of the German Federal Sate of Lower Saxony (Niedersächsisches Landesamt für Verbraucherschutz und Umwelt, LAVES). *ACOD1^-/-^*, *ACOD1^+/-^,* and *ACOD1^+/+^* mice with C57BL/6J background had originally been generated by Dr. Haruhiko Koseki in the RIKEN Institute (Yokohama, Japan) by the use of stem cells which were purchased from the Repository of Knockout Mouse Project under strain ID Irg1tm1a (KOMP) Wtsi. Age-matched groups of female *ACOD1^-/-^, ACOD1^+/-^*, and *ACOD1^+/+^* mice were derived from the same heterozygous breeding pair. Animals were bred at the University of Luxembourg and then transferred to the Animal Experimental Unit of the Helmholtz-Centre for Infection Research, where they were kept under Animal Biosafety Level 2 and non-specific pathogen-free conditions. Animals were housed seperated according to genotype, up to 5 mice per cage. For IAV infections, mice were sedated by intraperitoneal injection of ketamine (10 mg/mL) and xylazine (1 mg/mL) in 0.9% NaCl, infected intranasally and then observed for weight loss and survival for up to 15 days. Mice with weight loss >20% had to be sacrificed (CO_2_ asphyxiation) and were counted as non-survivors. In a separate experiment, mice were sacrificed on days 8/9 and 14 p.i. (one mouse that died on day 11 was also included) and lungs examined for histopathological changes. Formalin-fixed, paraffin-embedded sections of mouse lungs were stained with hematoxylin/eosin (H&E). To assess the degree of inflammation and histopathological lesions, 10 parameters of inflammation and tissue damage were scored by an expert in mouse pathology (MP), who was not aware of the identity of the slides, according to the scale below.

### Human lung tissue explant model

The study was approved by the Ethics Committee of Hannover Medical School (MHH), and all donors gave informed consent for experimental use of their tissue. Tissue was obtained at the time of medically indicated lung transplantation from patients with end-stage lung disease due to emphysema or pulmonary arterial hypertension. The overall tissue quality and gross pathological changes were assessed by a board-certified pathologist (Dept. of Pathology, Hannover Medical School). Healthy appearing tissue was then dissected and sectioned into pieces of approx. 30–50 mm^3^ (typical weight, 30-50 mg). Tissue pieces were cultured overnight in individual wells in RPMI medium, containing DI (1 mM) or itaconate (25 mM) as indicated. After this pre-treatment + washing step in medium, tissue pieces were infected with IAV (2×10^5^ FFU/ml) for 24 h at 37°C in medium containing DI or IA.

### RT-qPCR

RNA was purified using the Nucleospin RNA purification kit (Machery Nagel) and on-column removal of DNA by digestion with rDNase (Machery Nagel) for 15 min. at RT. cDNA was synthesized with the PrimeScript™ kit (TaKaRa, Shiga, Japan) using 400 ng RNA in a 10 µl reaction. RT-qPCR reactions were set up in a final volume of 20 µl, using the SensiFast™ SYBR® No-ROX Kit (Bioline, Taunton, MA) and the primers listed in Table 2. RT-qPCR was performed in a LightCycler® 2.0 instrument (Roche, Mannheim, Germany), using 45 cycles of the following program: 95°C for 15 sec., 60°C for 15 sec., and 72°C for 15 sec. To exclude artefacts resulting from primer dimer formation, melting curve analysis was performed using the sequence 95°C for 15 sec., 60°C for 15 sec., 95°C for 1 min. and 37°C for 30 sec. Relative expression of the host mRNA targets was calculated using the 2^−ΔΔCT^ method(Paijo, Döring et al., 2016), and expression of IAV *HA* mRNA by normalizing against expression of HPRT as internal control.

### Microarray analyses

RNA was extracted using the RNeasy kit (Qiagen), RNA quality was checked with a Bioanalyzer 2100 (Agilent Technologies) and samples with RNA integrity number >7 were used for microarray analysis. Total RNA/Poly-A RNA Control Mixture was prepared by adding poly-A RNA controls (Affymetrix). This poly-A RNA was then used to prepare double standard cDNA. Antisense RNA (complimentary RNA or cRNA) was then synthesized and amplified by *in vitro* transcription (IVT) of the ds-cDNA template using T7 RNA polymerase. Enzymes, salts, inorganic phosphates, and unincorporated nucleotides were removed to prepare the cRNA for 2nd-cycle ds-cDNA synthesis. After verifying quality and yield of cRNA, sense-strand cDNA was synthesized by the reverse transcription of cRNA. cRNA template was hydrolyzed leaving ds-cDNA, which was then purified. cDNA was fragmented and labeled by terminal deoxynucleotidyl transferase (TdT) using the Affymetrix proprietary DNA Labeling Reagent that is covalently linked to biotin. Cartridge Array Hybridization was conducted on the GeneChip® Instrument (Affymetrix) using Thermo Fisher Scientific microarrays (Clariom™ S Assay, human). Raw .CEL files from Clariom S Pico Assay microarrays for hg.38 were imported into the Transcriptome Analysis Console (TAC4.0.2) Software (ThermoFisher Scientific). Array QC and data normalization was performed using the robust multiarray average (RMA) method(Irizarry, Hobbs et al., 2003). Differentially expressed genes (DEGs) were identified using limma:Linear Models for microarray and RNA-Seq Data (www.Bioconductor.org) and one-way ANOVA.

### Differential gene expression analysis of bulk RNAseq data

Differential gene expression for Fig. 1B was calculated using generalized linear models contained in the edgeR package (doi:10.1093/bioinformatics/btp616). P values were adjusted using false discovery rate (FDR), using FDR <0.05 to define significance of differential gene expression.

### Single cell RNA sequencing

#### Barcoding and cDNA synthesis

The single cell suspension was loaded onto a well on a 10x Chromium Single Cell instrument (10x Genomics). Barcoding and cDNA synthesis were performed according to the manufacturer’s instructions. Briefly, the 10x™ GemCode™ Technology partitions thousands of cells into nanoliter-scale Gel Bead-In-EMulsions (GEMs), where all the cDNA generated from an individual cell share a common 10x Barcode. In order to identify the PCR duplicates, Unique Molecular Identifier (UMI) was also added. The GEMs were incubated with enzymes to produce full length cDNA, which was then amplified by PCR to generate enough quantity for library construction. Quality was checked using the Agilent Bioanalyzer High Sensitivity Assay.

#### Library construction and quality control

The cDNA libraries were constructed using the 10x ChromiumTM Single cell 3’ Library Kit according to the manufacturer’s original protocol. Briefly, after the cDNA amplification, enzymatic fragmentation and size selection were performed using SPRI select reagent (Beckman Coulter, Cat# B23317) to optimize the cDNA size. P5, P7, a sample index and read 2 (R2) primer sequence were added by end repair, A-tailing, adaptor ligation and sample-index PCR. The final single cell 3’ library contains a standard Illumina paired-end constructs (P5 and P7), Read 1 (R1) primer sequence, 16 bp 10x barcode, 10 bp randomer, 89 bp cDNA fragments, R2 primer sequence and 8 bp sample index. For post library construction QC, 1 μl of sample was diluted 1:10 and ran on the Agilent Bioanalyzer High Sensitivity chip for qualitative analysis.

#### Single-cell RNA sequencing and generation of data matrix

PBMC were infected and treated as for the microarray analyses. Cells were then stained with propidium iodide (Apoptosis Detection Kit, eBioscience, cat. no. 88-8005-74), and dead cells and doublets were removed by FACS. Live cells (including those undergoing apoptosis) were used to make single-cell suspensions using the 10x Genomics platform. Libraries were sequenced on an Illumina NovaSeq 6000 2×50 paired-end kits using the following read length: 26 bp Read1 for cell barcode and UMI, 8 bp I7 index for sample index and 89 bp Read2 for transcript. Cell Ranger 1.3 (http://10xgenomics.com) was used to process Chromium single-cell 3’ RNA-seq output. First, “cellranger mkfastq” demultiplexed the sequencing samples based on the 8bp sample index read to generate fastq files for the Read1 and Read2, followed by extraction of 16bp cell barcode and 10bp UMI. Second, “cellranger count” aligned the Read2 to the pre-built human reference genome (hg19, GRCh38) using STAR. Then, aligned reads were used to generate data matrix only when they had valid barcodes and UMI, and mapped to exons (Ensembl GTFs GRCm38.p4) without PCR duplicates. Valid cell barcodes were defined based on UMI distribution.

#### Single-cell gene expression data analysis including filtering, normalization, and clustering

was processed using Seurat V3.1 (https://satijalab.org/seurat/) in the R environment (www.bioconductor). Cells were removed when they had < 500 genes and > 25% reads mapped to mitochondrial expression genome. After filtering and doublet removal, 6456 single cells (control = 1870, DI =1474, IAV=1701, IAV & DI= 1411) remained. Raw counts were normalized by the global-scaling normalization method “LogNormalize”. Principal component analysis score was used to determine the ‘dimensionality’ of all cells. Scatter plots were obtained using the UMAP method. Cell clusters were identified as different cell populations based on the expression of recently published canonical markers (Table 3). Differential gene expression was determined for each cell type using DESeq2 (Love, Huber et al., 2014). IAV infected cells were identified by the presence of IAV encoded transcripts, as mapped against NIAID Influenza Research Database (Zhang, Aevermann et al., 2017), www.fludb.org/brc/fluStrainDetails.spg?strainName=A/Giessen/6/2009(H1N1)&decorator=influenza . mRNA encoding the viral M protein could not be identified, likely due to a sequence mismatch.

### Enzyme-linked immunoassay

Supernatants from the same PBMC as used for RNA analyses were collected 12 h p.i. Protein concentrations were measured using ELISA MAX Deluxe Sets (Biolegend; IL1B cat. no. 437004; IL6, 430501; TNFa, 430201; IFNG, 430101) as per the manufacturer’s instructions. Briefly, capture antibody was used to pre-coat the wells O/N. After removing the capture antibody and washing the plates, block buffer was applied for 1 h at room temperature at 500 rpm. Again, plates were washed at least 4 times before adding the samples (100 μL, diluted as needed) and standards for 2 h at RT. The plates were then washed again 4 times, and detection antibody was added to each well. The plate was further incubated for 1 h at RT and then washed 4 times. Avidin-HRP conjugate was added and incubated for 30 min. Substrate was added until color developed, and the samples were measured after 15-20 min at 450 and 570 nm.

### Multiplex assay for inflammation-related polypeptides

Concentration of 27 inflammation-related human proteins were measured in cell culture supernatants by using Human 27-plex BIORAD cytokine panel (Cat. No.171-A1112, BIORAD). Standard curves were generated for all 27 cytokine standards with eight 2-fold serial dilutions starting from 32 ng/ml. 100 µl of assay buffer was added to each well of a microfiltration plate, followed by addition of 50 µl of beads suspension. After washing the beads with assay buffer, 50 µl of standard or the sample (supernatant) was added to each well and incubated for 30 min at room temperature with gentle shaking. 25 µl of antibody premix for detection was added into each well followed by incubation for 30 min at room temperature with gentle shaking. Three washing steps were performed using 50 µl of assay buffer in each well, followed by washing with streptavidin solution for 10 min at room temperature with shaking. Final washing with 125 µl of assay buffer was performed before quantification by using BIO-PLEX Manager software version 4.

### Immunofluorescense

THP-1 cells were seeded and differentiated with 80% confluency in a 24 well plate with coverslips. Cells were fixed with 2.5% formaldehyde for 20 min and the reaction stopped by adding 0.1 M glycine for 5 minutes (min). Cells were then permeabilized with 0.2% Triton ×100 in PBS for 5 min. Cells were incubated for 1 h with primary antibody (rabbit-anti p65 mAB, Cell Signaling, cat. no. 8242S) 1:200 diluted in BSA/PBS 1 mg/ml. Cells were then washed three times with PBS and incubated in the dark for 45 min with Alexafluor594 secondary antibody (1:100) (Cell Signaling, cat.# 8889S) in BSA/PBS 1 mg/ml and incubated in the dark for 45 min. Cells were washed three times with PBS. To stain nuclei, DAPI was added at 1:10,000 in PBS for 10 min. Cells were washed three times with PBS and mounted with 20 μl of DAKO mounting medium per coverslip on glass slides and stored in the dark until imaged. The images were analysed using Image J s/w (imagej.net).

### Quantification of itaconate

Itaconate concentrations were measured by liquid-chromatography tandem mass-spectrometry (HPLC-MS/MS) using a liquid chromatography system (NEXERA LC, Shimadzu, Japan) coupled to a triple quadrupole - ion trap mass spectrometer (QTRAP 5500, Sciex, Framingham, MA) and Analyst 1.7 software (Sciex, Framingham, MA, USA), as described in detail in (Kuhn, 2017).

### Statistics

Data are shown mean ± standard error of the mean (SEM) unless stated otherwise. Significance of between-group and across-group differences was assessed with the statistical tests indicated in the figure legends, using Prism 5.02 and 8.0 (GraphPad Software Inc.). Wherever exact p values are not stated, significance is graded as *p<0.05, **p<0.01, ***p<0.001.

### Principle component analysis

was performed with the R package “PCAtools” (https/github.com/kevinblighe/PCAtools), and **hierarchical clustering analysis** with the R tool pheatmap (cran.r-project.org/web/packages/pheatmap/pheatmap.pdf).

### Reanalysis of published data sets

Reanalyses of publically available published data sets were performed in the R environment (version 3.6.3) (R_Core_Team, 2014), using package beeswarm (version 0.2.3.) (Eklund, 2016) for visualization as indicated.

### Gene set enrichment analysis

Features with linear fold change (FC) ≥|1.5| and unadjusted P-value ≤0.05 were used for enrichment analysis (Subramanian, Tamayo et al., 2005) based on gene ontology (GO Biological Process) and KEGG pathways, using GEne SeT AnaLysis Toolkit (Liao, Wang et al., 2019). Enrichment/depletion strip charts were made using ggplot2 (Wickham, 2016). Volcano plots to shows DEGs were made using R package “EnhancedVolcano” (https://github.com/kevinblighe/EnhancedVolcano). Molecular Signatures Database (MSigDB) v7.2 (Liberzon, Birger et al., 2015) was used to collect the information of genes involved in gene set enrichment analyses.

## Supplemental figure legends

**Figure S1.**
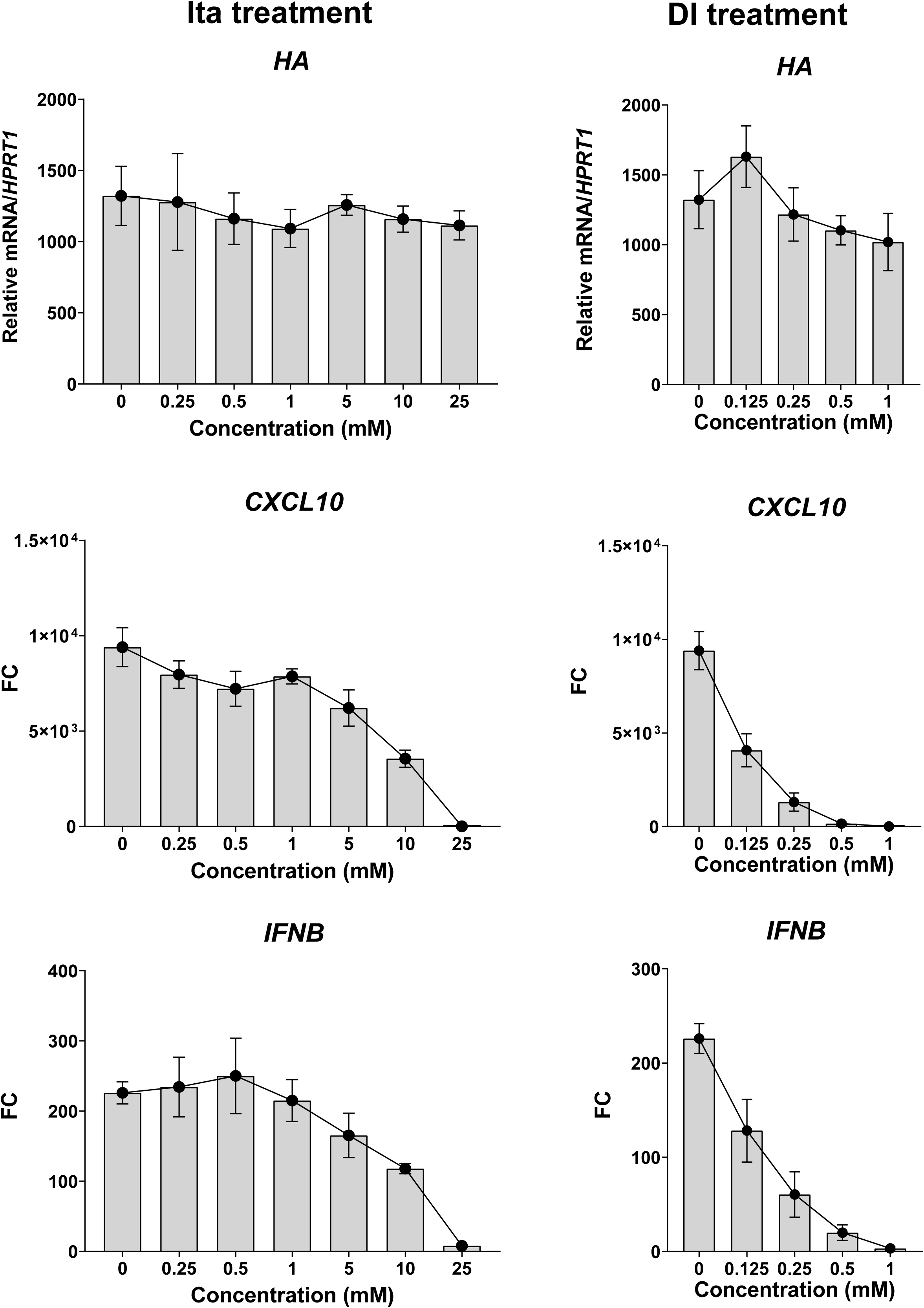
Dose response curves of itaconate and DI treatment of IAV-infected dTHP-1 cells. dTHP-1 cells were pretreated overnight with the indicated concentrations of IA or DI, infected with IAV, and expression of *HA* and *CXCL10* mRNA was measured after 12 h (RT-qPCR) (n=3).

**Figure S2.**
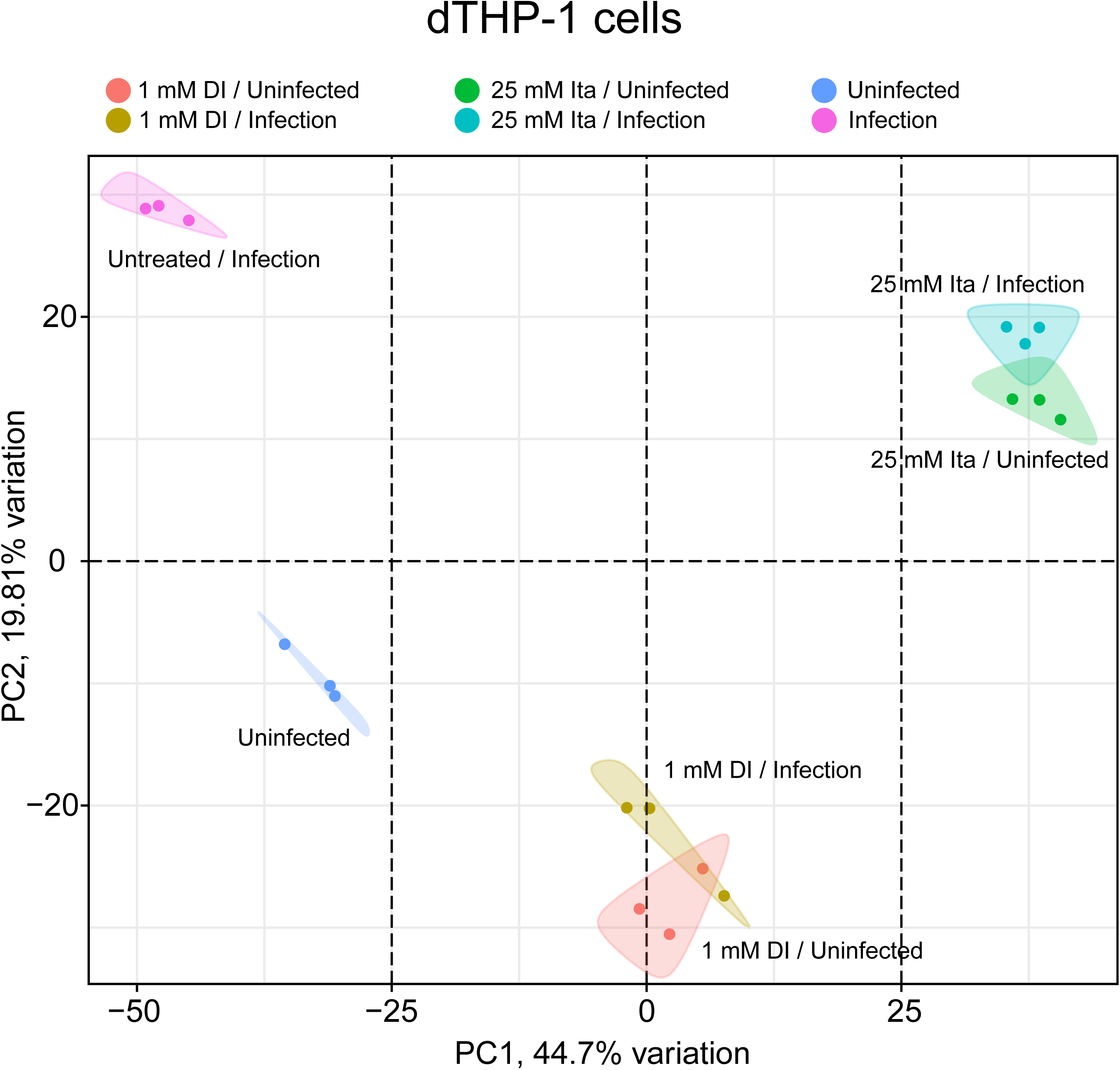
The impact of itaconate and DI treatments on cell transcriptomes is greater than that of IAV infection. PCA based on the microarray analysis shown in Fig. 3. Untreated and treated IAV-infected cells cluster together, whereas untreated infected cells are clearly separated from all other groups in both principle components.

**Figure S3.**
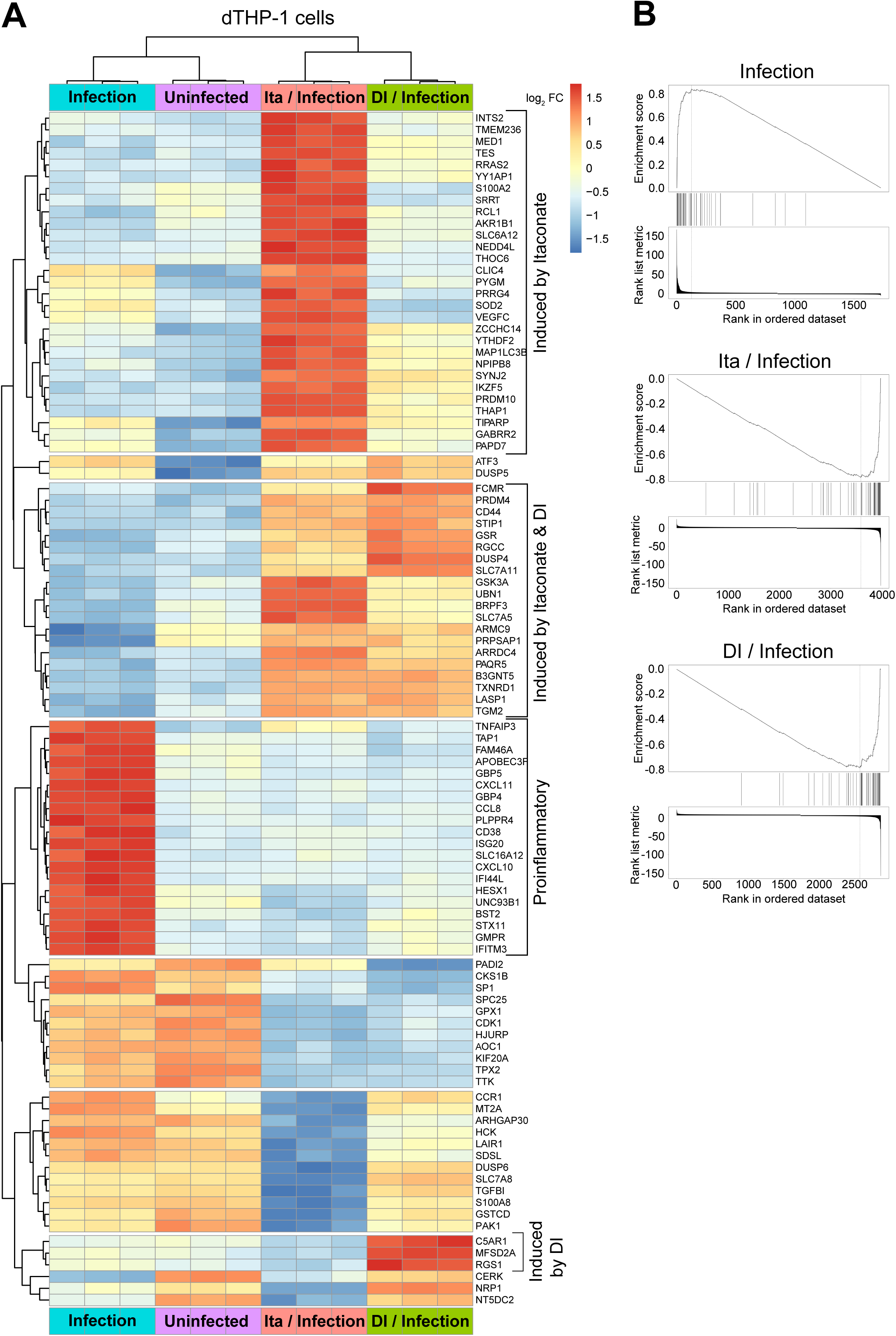
A. Hierarchical clustering analysis of effects of itaconate and DI on IAV-infected dTHP-1 cells. Analysis based on the same microarray data as used for Fig. 3 and EV1B, using the top 100 DEGs (FDR F-test <1.89E-09). Uninfected cells and untreated IAV-infected cells form one clade and itaconate and DI treated IAV-infected cells the other clade. The brackets identify (i) a large pro-inflammatory subclade in IAV infection, which is globally downregulated by the treatments; (ii) a large cluster of DEGs specific for itaconate treatment; (iii) a small cluster of DEGs specific for DI treatment, and (iv) a subclade comprising DEGs differentially expressed due to both compounds. **B. Enrichment plots based on the 99 genes contained in GO term *Response to type I interferon*.** Enrichment scores are plotted on the y-axis, the mRNAs (identified by vertical lines) along the x-axis, ranked by fold change. The plots illustrate pronounced enrichment in a large number of genes due to infection, which is nearly quantitatively depleted by treatment with both itaconate and DI.

**Figure S4.**
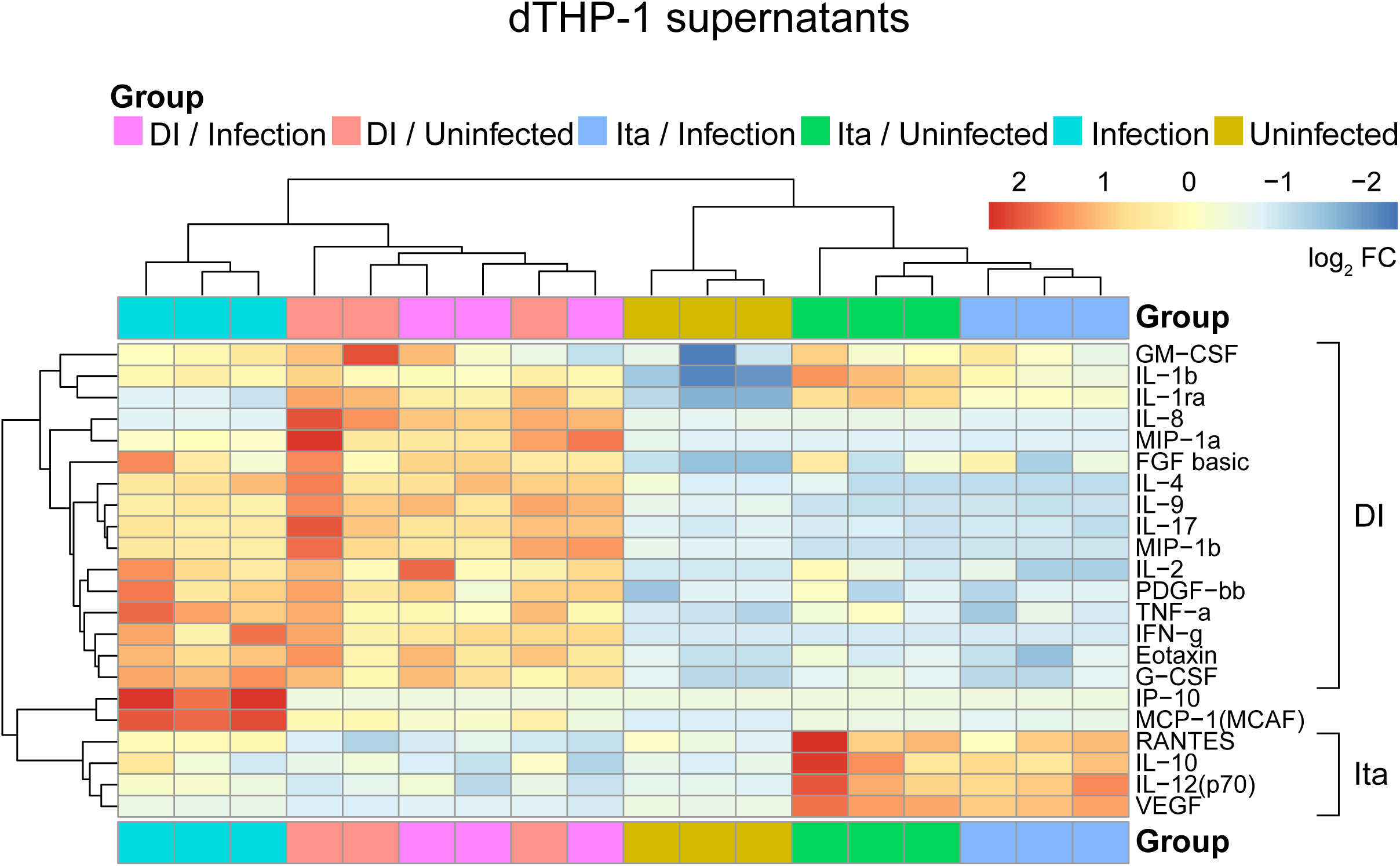
Itaconate and DI induce different recruitment programs in dTHP1 cells. Supernatants of uninfected and IAV-infected dTHP1 cells, with our without itaconate or DI treatment, were analyzed for concentrations of 27 cytokines/chemokines by multiplex microbead array 12 h after infection or mock treatment. A hierarchical clustering analysis was carried out with those targets that were detected above limit of detection of the assay in most samples. The microarray analysis of cellular gene expression from the same experiment is shown in Fig. 4 (n=3).

**Figure S5.**
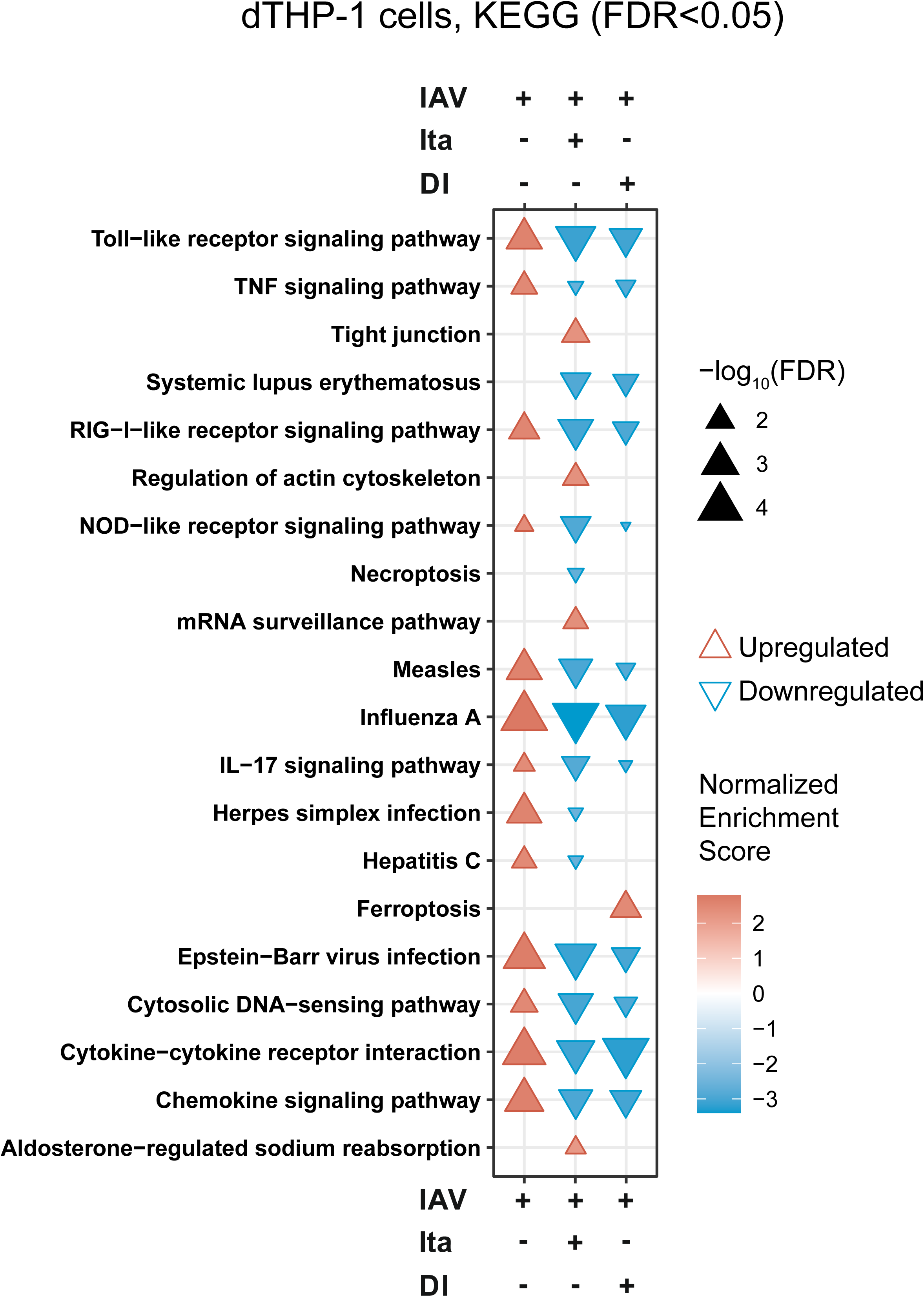
KEGG-pathway analysis of effects of itaconate and DI treatment on IAV-infected dTHP-1 cells. Analysis based on the microarray data set used for Figure 4. A broad dampening of IFN-related and pro-inflammatory pathways by both compounds is evident.

**Figure S6.**
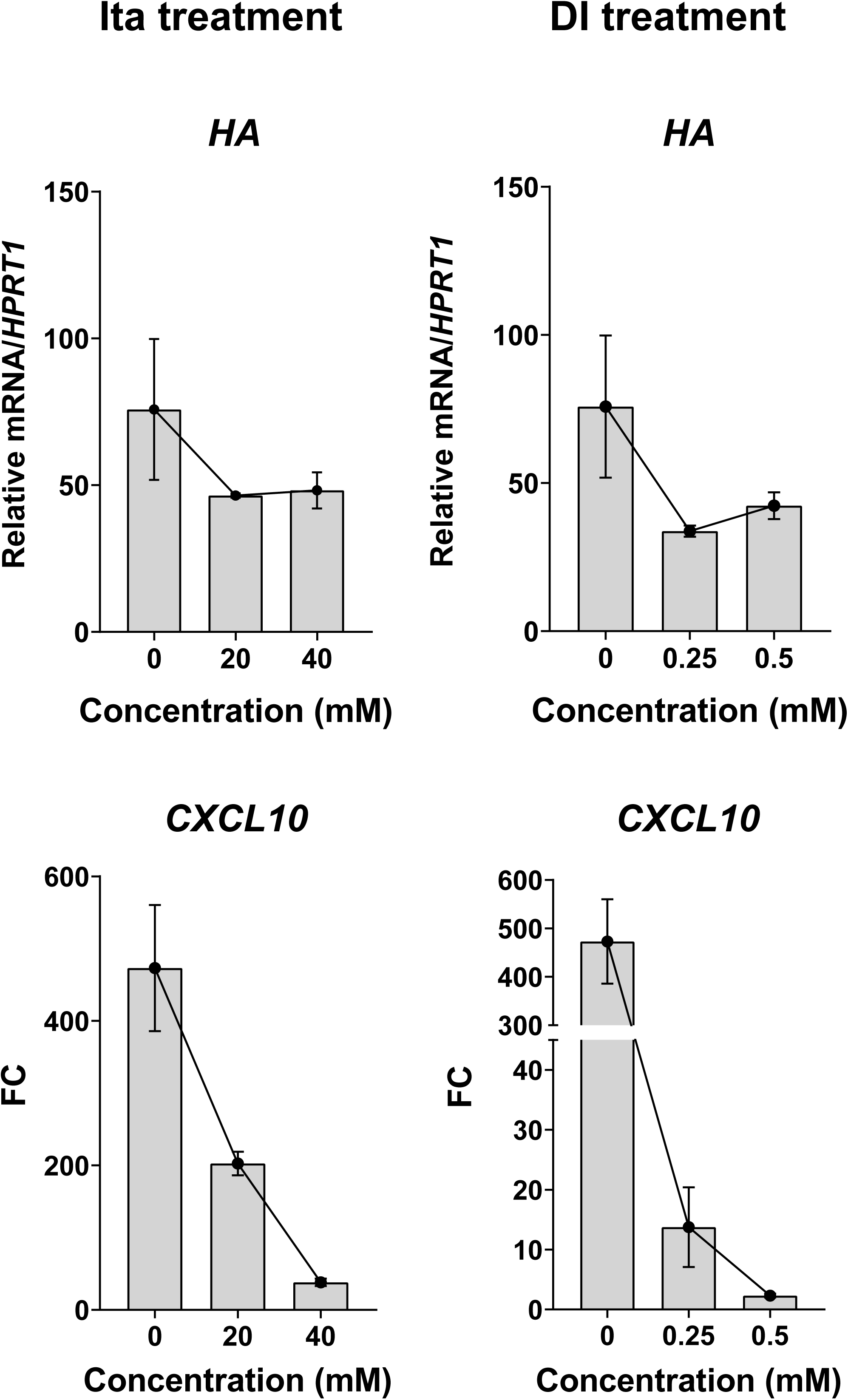
Effects of increasing doses of itaconate and DI on *CXCL10* expression in IAV-infected A549 cells. RT-qPCR using *HPRT1* as reference. A549 cells were infected with IAV and itaconate and DI treatments were given at the indicated concentrations. Expression of *HA* and *CXCL10* mRNA was measured by RT-qPCR 24 h p.i. (n=3). DI concentrations ≥ 1 mM could not be evaluated due to widespread cytotoxicity seen by light microscopy.

**Figure S7.**
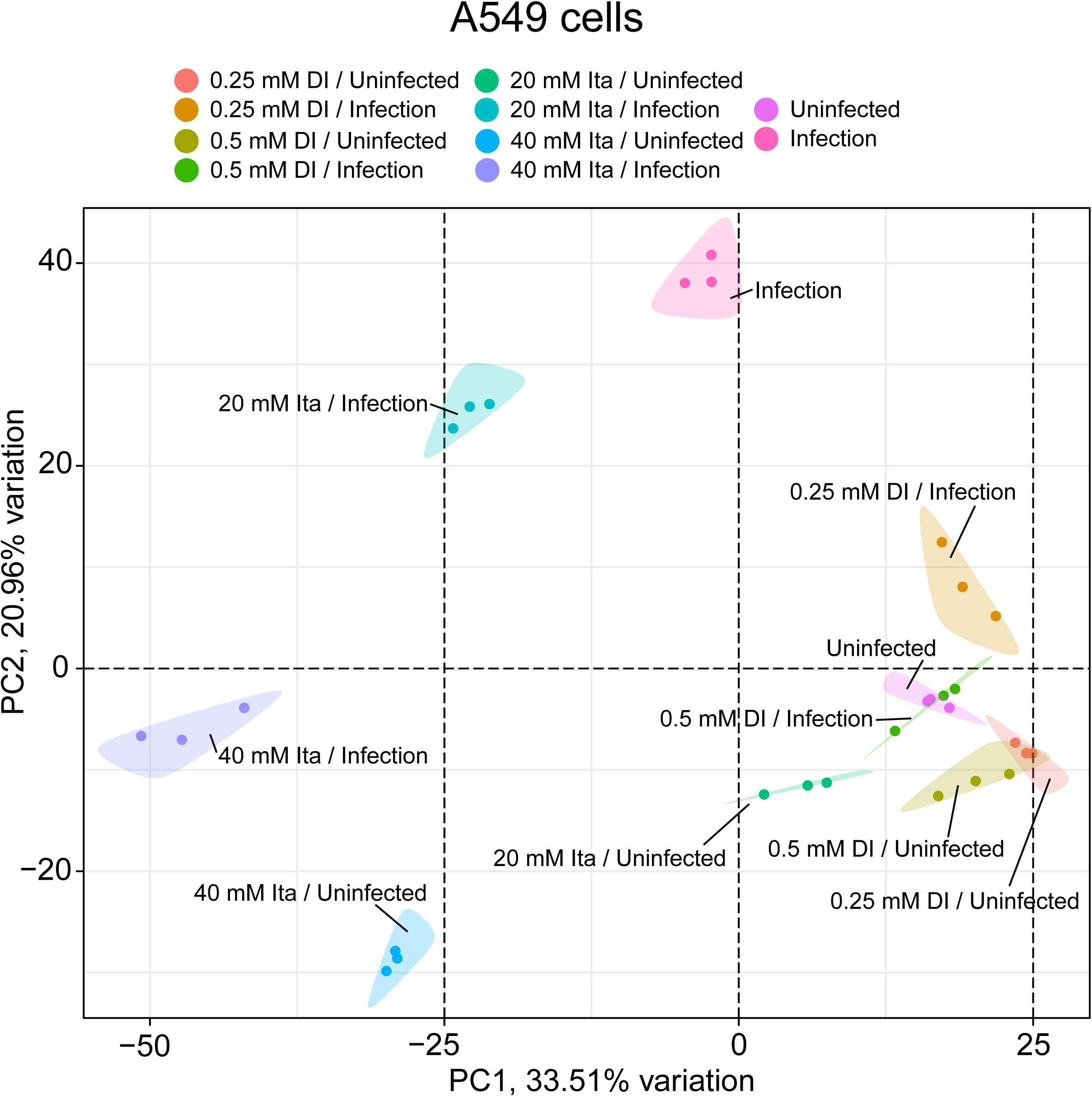
Global effects of itaconate and DI on transcriptomic responses in IAV-infected A549 cells. PCA based on the microarray analysis used for Fig. 6, but additionally including treatments with 40 mM itaconate and 0.25 mM DI. PCA showing normalization of IAV-driven reprogramming of gene expression, but increasing impact of itaconate on cellular responses with increasing doses.

**Figure S8.**
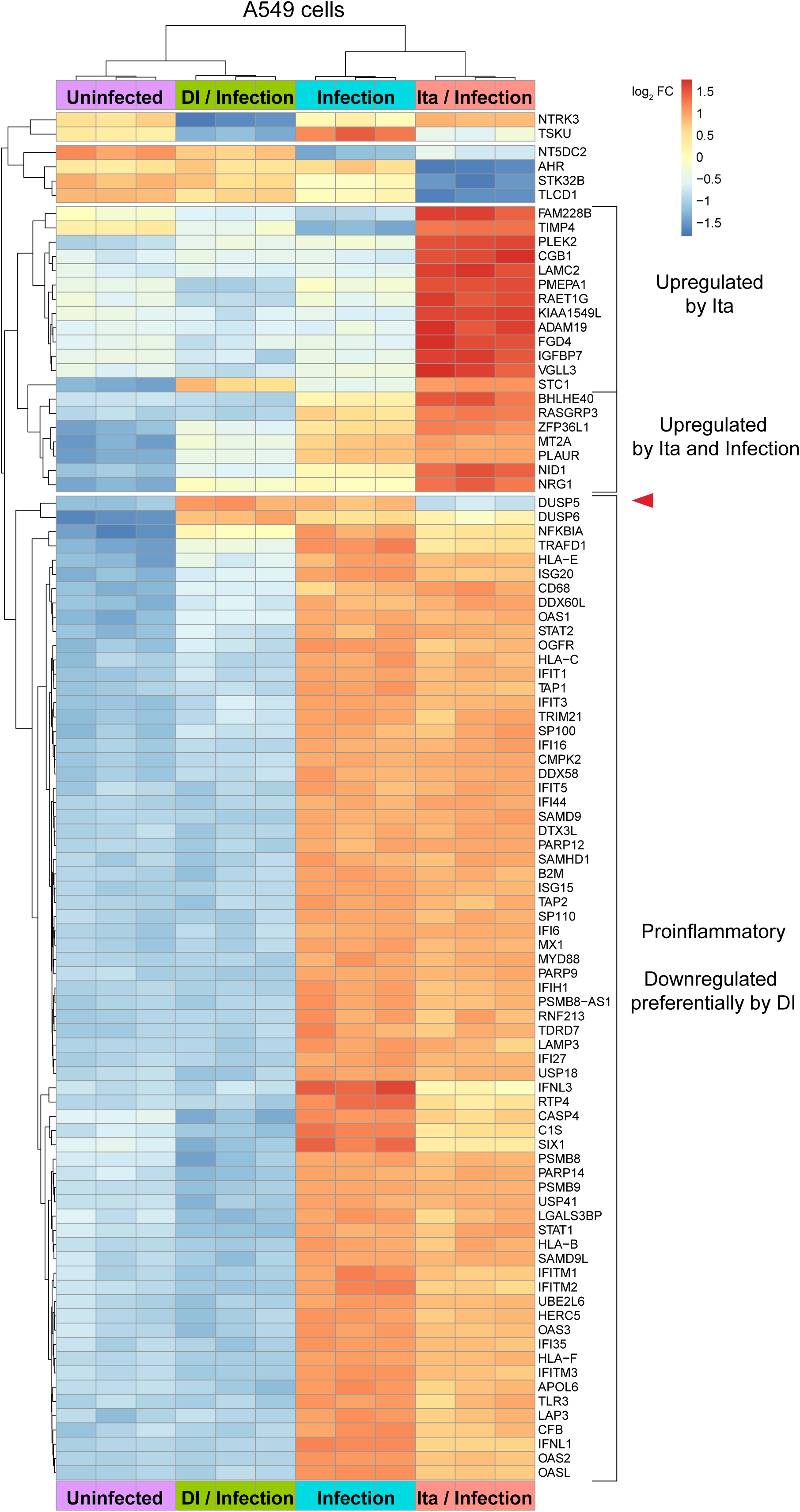
Effects of itaconate (20 mM) and DI (0.5 mM) on transcriptomic responses in IAV-infected A549 cells. Analysis based on the microarray data used for the GO analysis in Fig. 6. Unsupervised hierarchical clustering analysis of the 100 most significant DEGs (FDR F-test <1.46E-10). DI-treated infected and uninfected cells cluster in one clade, and infected and itaconate-treated infected cells in the other. There is a large clade mostly containing proinflammatory genes, which are downregulated by DI, whereas effects of itaconate are much weaker. *DUSP95* is exclusively downregulated by itaconate (arrowhead). There are two smaller clades of genes that form an “itaconate signature”, comprising genes that are, albeit to a lesser extent, also upregulated by IAV infection, or uniquely by itaconate.

**Figure S9.**
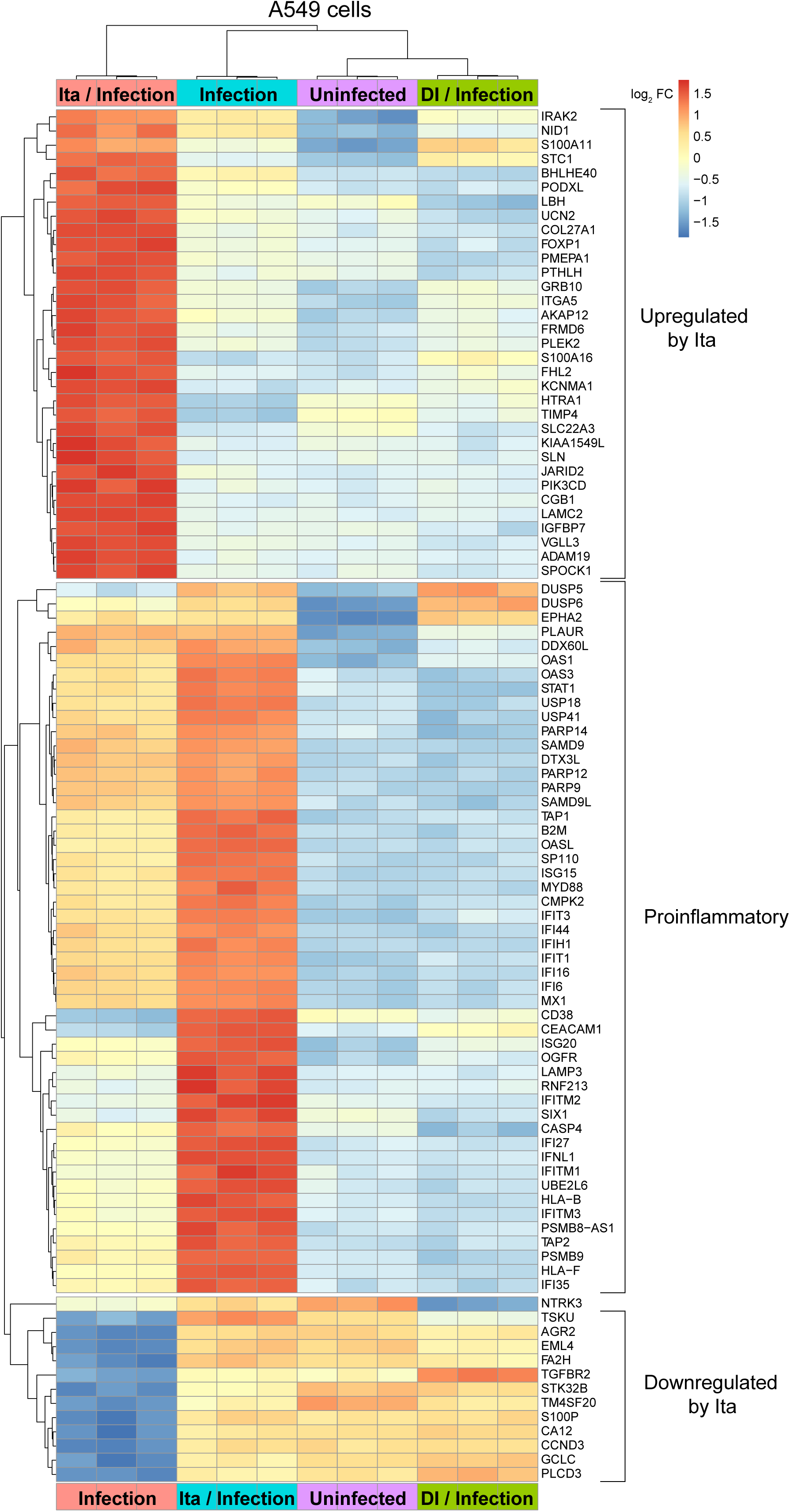
Effects of itaconate (40 mM) and DI (0.5 mM) on transcriptomic responses in IAV-infected A549 cells. Hierarchical clustering analysis ased on the microarray analysis used for Fig. S5. Compared to the 20 mM itaconate concentration (Fig. S8), itaconate-treated IAV infection is now in a separate clade, indicating that the impact of itaconate on the cells dominates that of the infection. Downregulation of the inflammation-driven clade is now more pronounced, the itaconate-unique signature is stronger, and there also is a clade comprised of genes downregulated by IAV.

**Figure S10.**
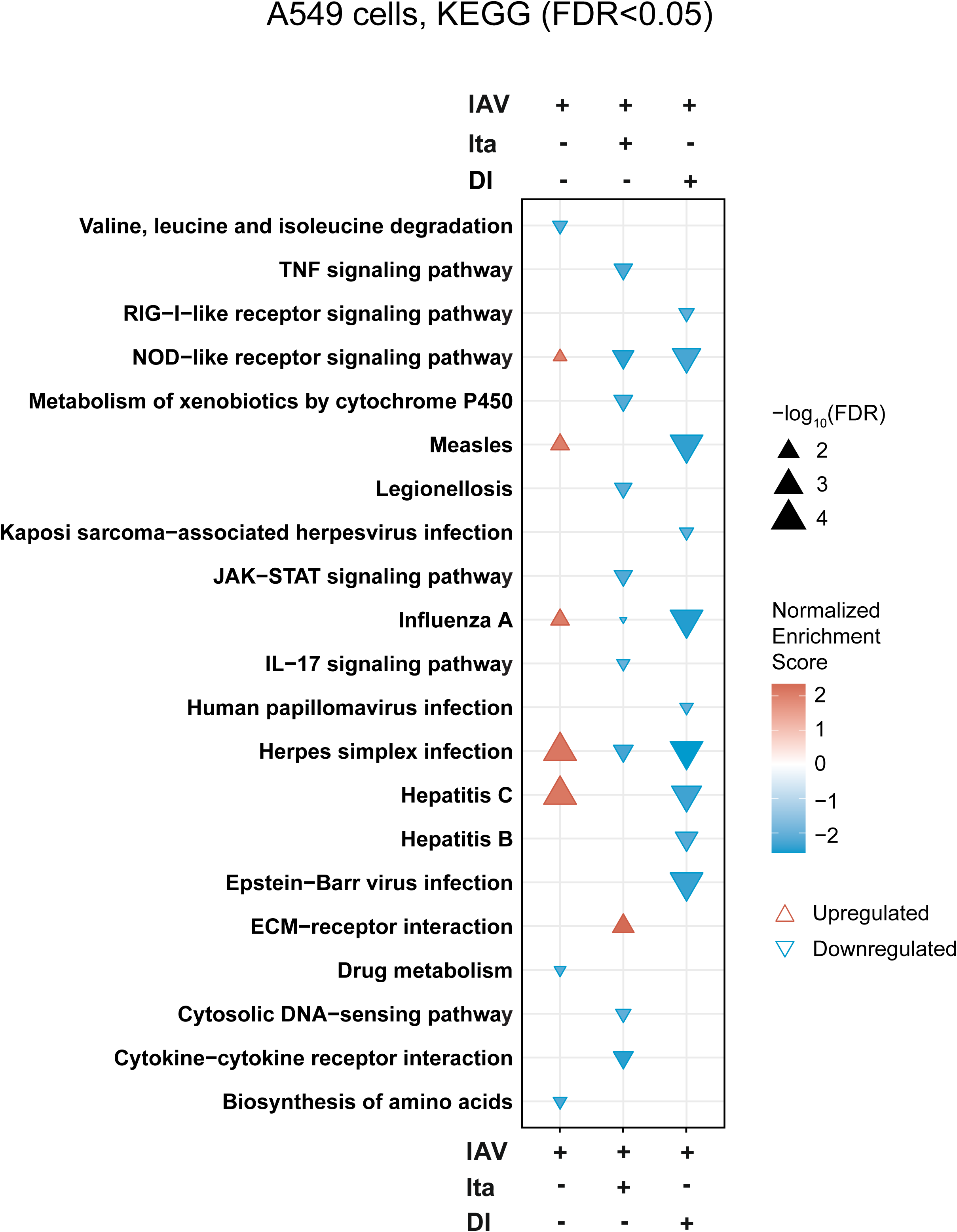
KEGG pathway enrichment analysis of itaconate (20 mM) and DI (0.5 mM) treatment on IAV infection in A549 cells. Analysis performed based on the microarray data shown in Fig. 6 and S8.

**Figure S11.**
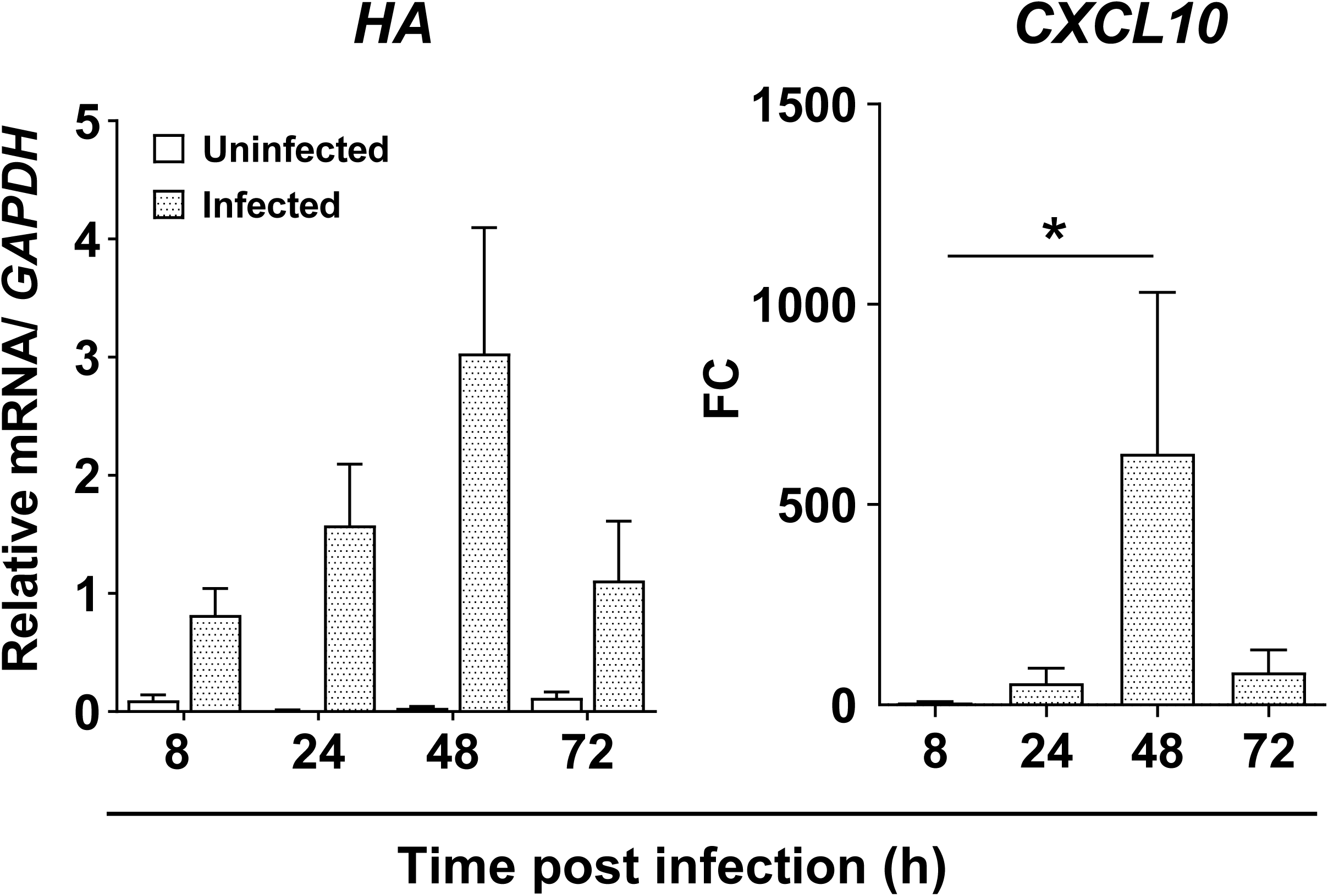
Primary human lung tissue explants support IAV RNA transcription and upregulation of CXCL10 mRNA. Human primary lung tissue from patients with emphysema or pulmonary arterial hypertension (n=7 donors, 3 tissue pieces per donor per treatment) was infected with IAV for 72 h and *HA* and *CXCL10* mRNA expression measured by RT-qPCR, using *HPRT1* as internal reference. *CXCL10* levels are expressed as fold change with reference to uninfected tissue at the same time point. *p<0.05; **p<0.01; ***p<0.001 (Mann-Whitney U test).

**Figure S12.**
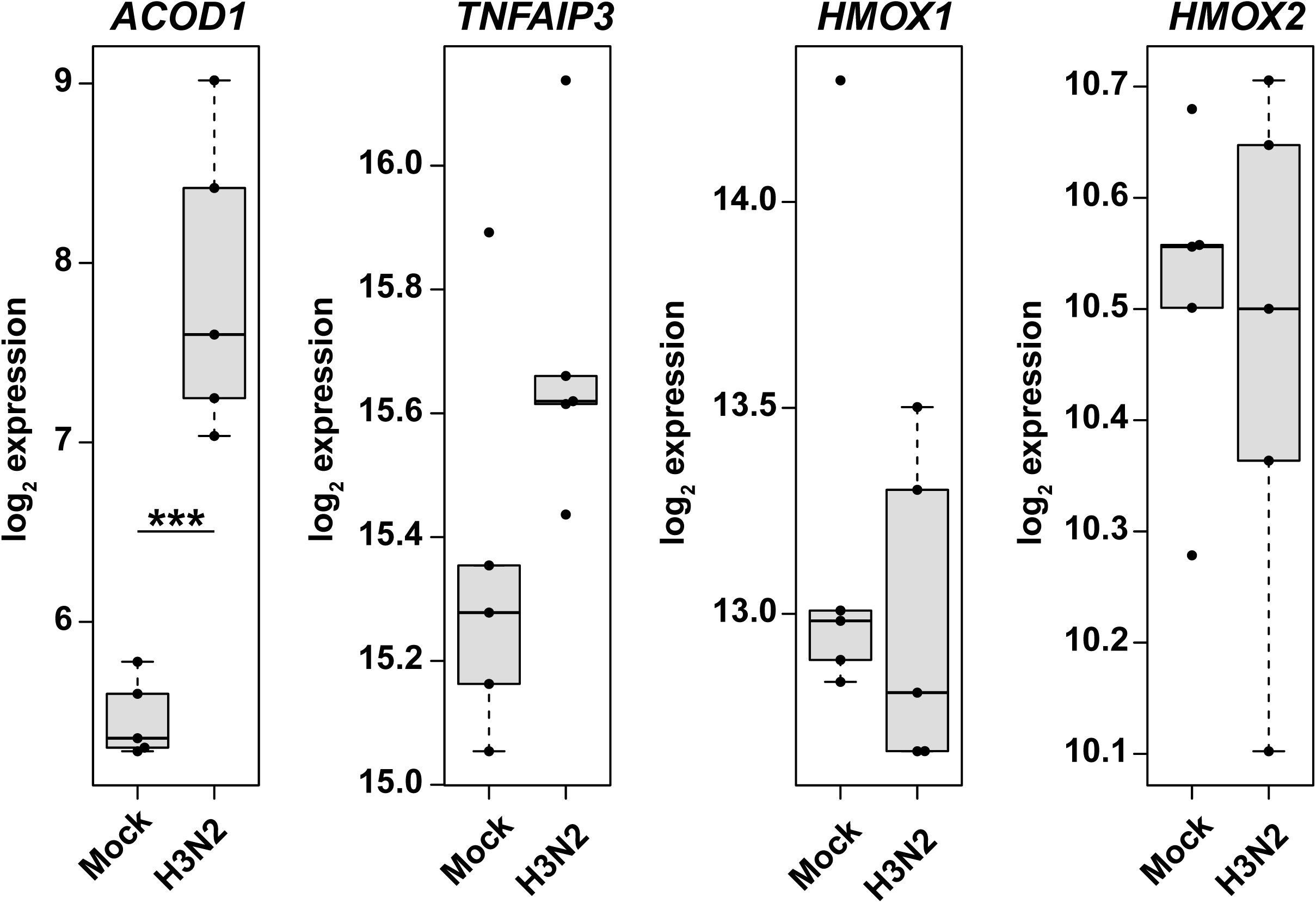
Induction of *ACOD1* and *TNFAIP3* mRNAs during IAV (H3N2) infection of human lung explants. Reanalysis of a published dataset of gene expression (RNAseq) in IAV (H3N2) infection of human lung tissue derived from tumor-free margins obtained during lobectomy for lung carcinoma(Matos et al., 2019). A strong induction of *ACOD1* expression and a tendency towards increased *TNFAIP3* expression are seen. As opposed to the strong induction of both genes in the mouse model (Fig. 1A), *HMOX1* and *HMOX2* expression is unchanged. *p<0.05; **p<0.01; ***p<0.001 (pairwise t-tests with pooled SD).

**Figure S13.**
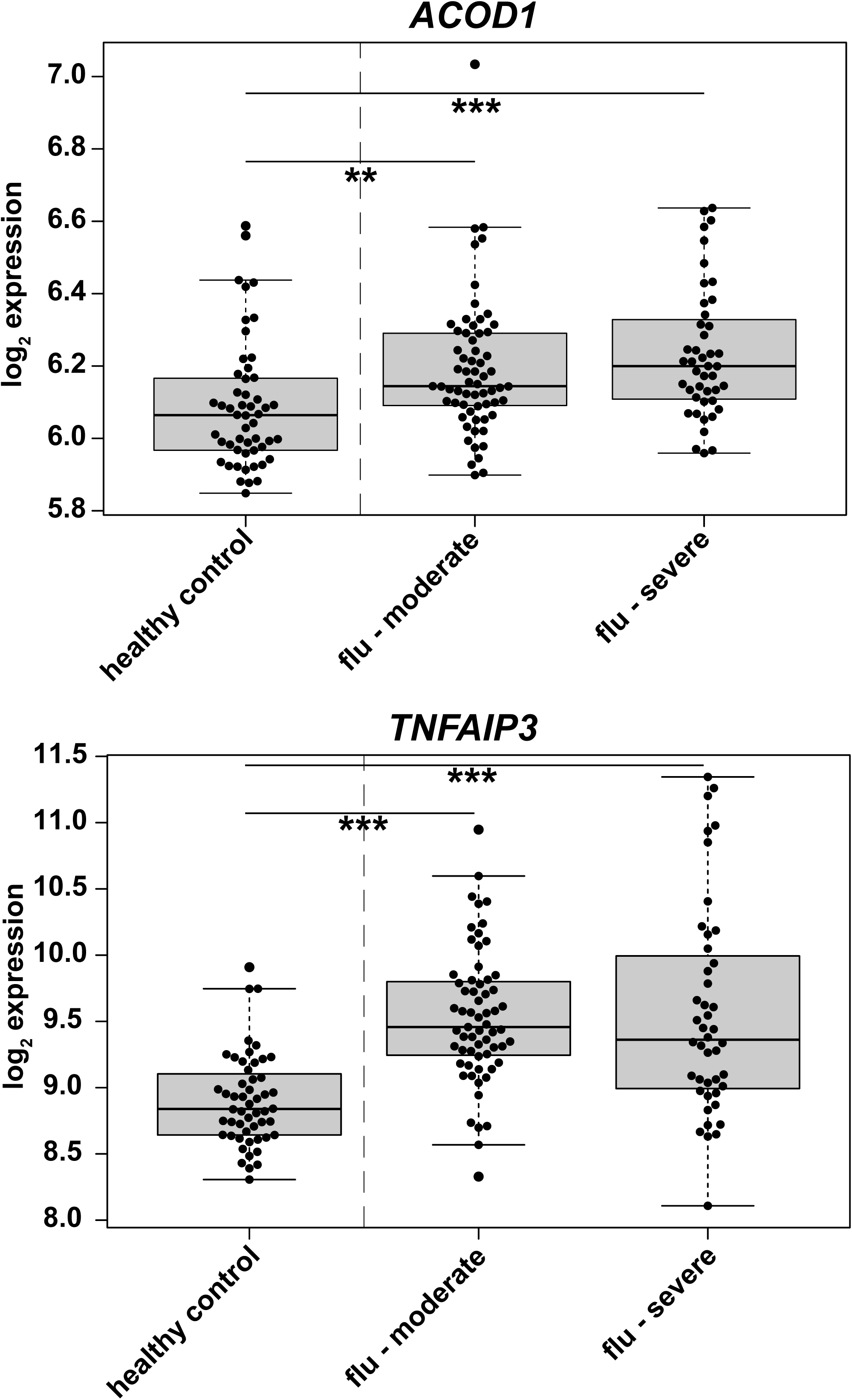
Increased levels of *ACOD1* and *TNFAIP3* mRNA expression in whole blood from patients with moderate and severe influenza. Reanalysis of a published data set of gene expression in whole blood from patients with moderate and severe influenza and healthy controls(Tang, Shojaei et al., 2017). *p<0.05; **p<0.01; ***p<0.001 (pairwise t-tests with pooled SD).

**Figure S14.**
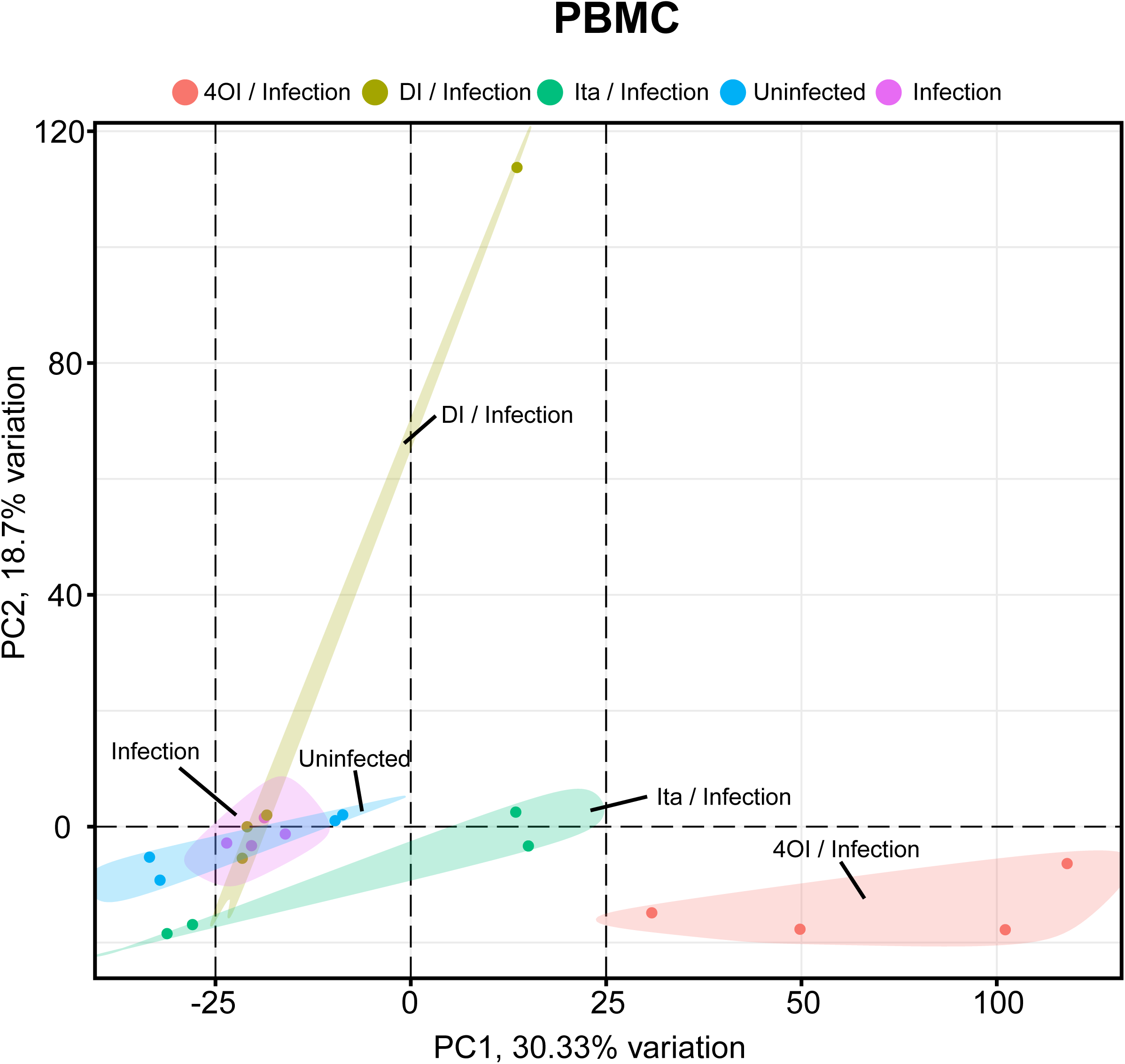
Transcriptome changes in itaconate, DI, and 4OI treated PBMC are driven more by the treatments than by infection. Analysis of mRNA expression in a subgroup (n=4) of the PBMC samples that were used for the targeted assays shown in Fig. 8. A PCA was performed on the same oligonucleotide microarray data as used for Fig. 8C and J. There is a less pronounced effect of IAV infection (control and IAV infected samples are not clearly separated) than on dTHP-1 and A549 cells, but pronounced additional broad changes in gene expression due to treatment with 4OI, itaconate treatment (note two outliers), and DI (one outlier).

**Figure S15.**
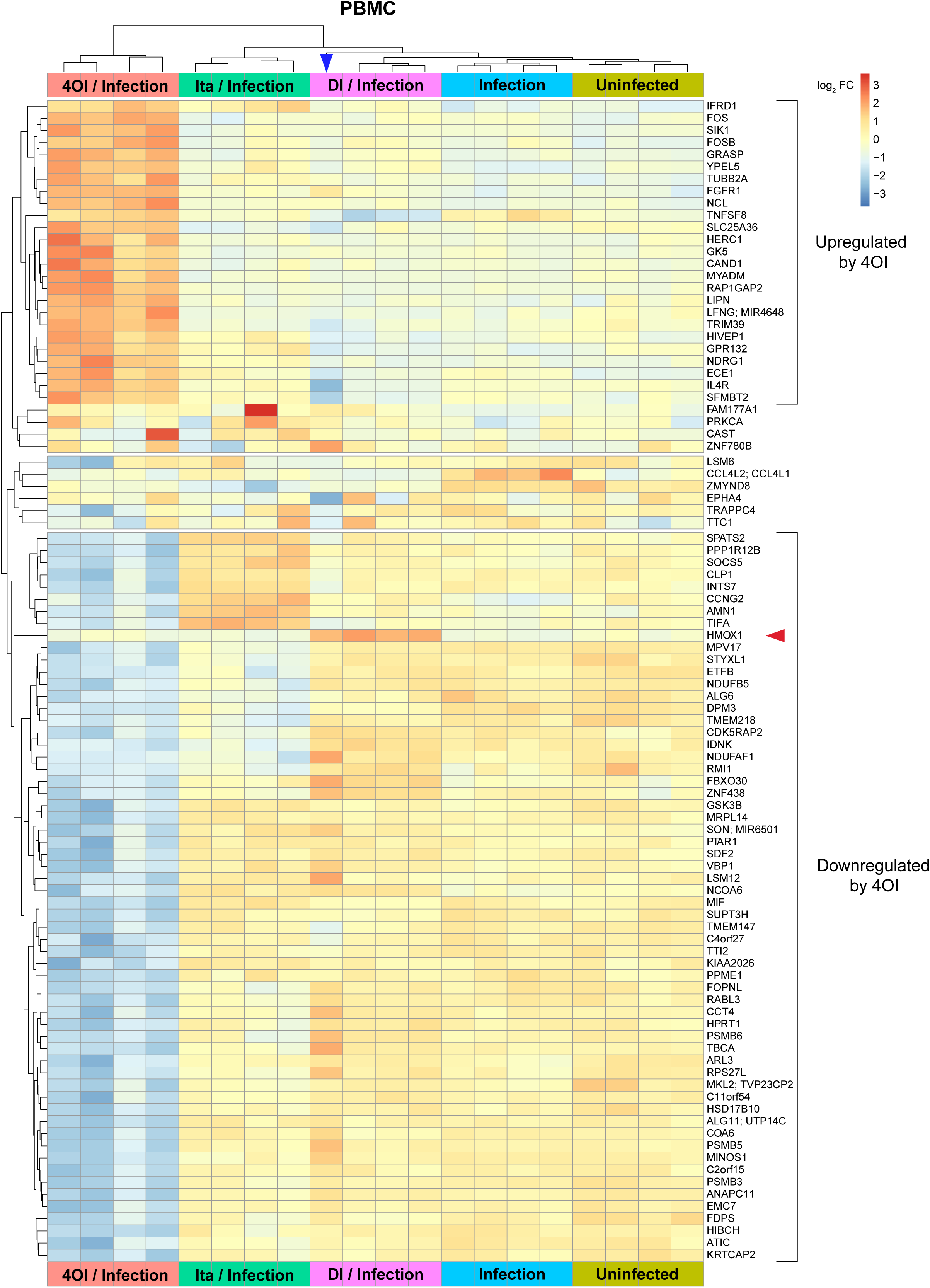
Hierarchical clustering analysis of transcriptomic changes in IAV infected PBMC and responses to itaconate, DI, and 4OI treatment. The 100 most significant DEGs were selected (FDR F-test <3.72E-05). Transcriptome changes are mostly due to marked effects of 4OI (more down-than upregulation) on genes that are not affected by IAV infection, indicating general effects on cell homeostasis. However, induction of *HMOX1* by DI in IAV infection is evident (red arrowhead). The blue arrowhead points to the extreme outlier under DI treatment seen in the PCA.

**Figure S16.**
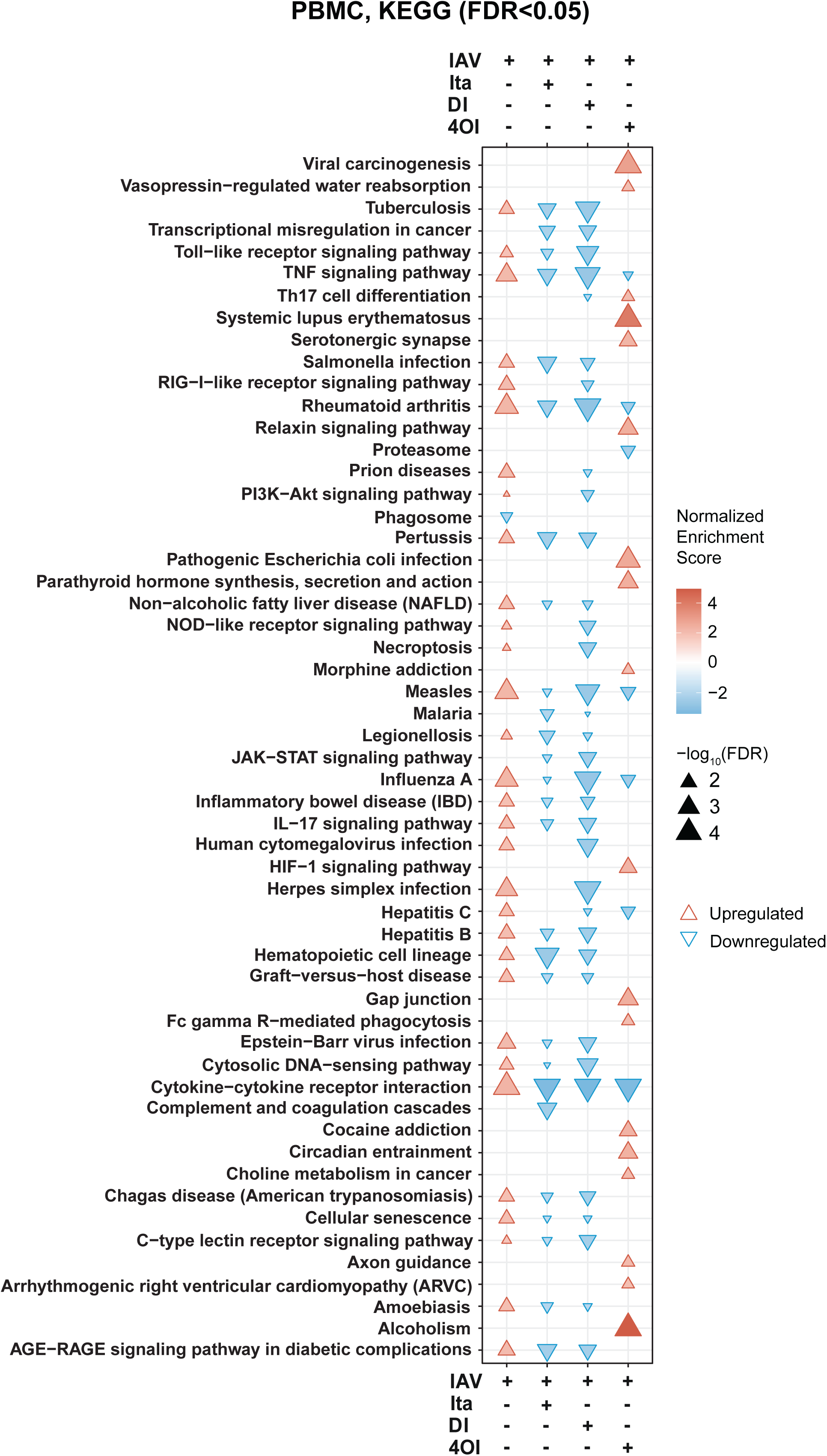
KEGG pathway analysis of transcriptomic changes in IAV infected PBMC and responses to itaconate, DI, and 4OI treatment. KEGG terms with an FDR <0.05 in at least one group were selected. The threshold for inclusion in the chart was raised to FDR <0.01 if a pathway was enriched/depleted only in a single treatment and not by IAV infection alone (e.g., *Alcoholism* in 4OI treatment).

**Figure S17.**
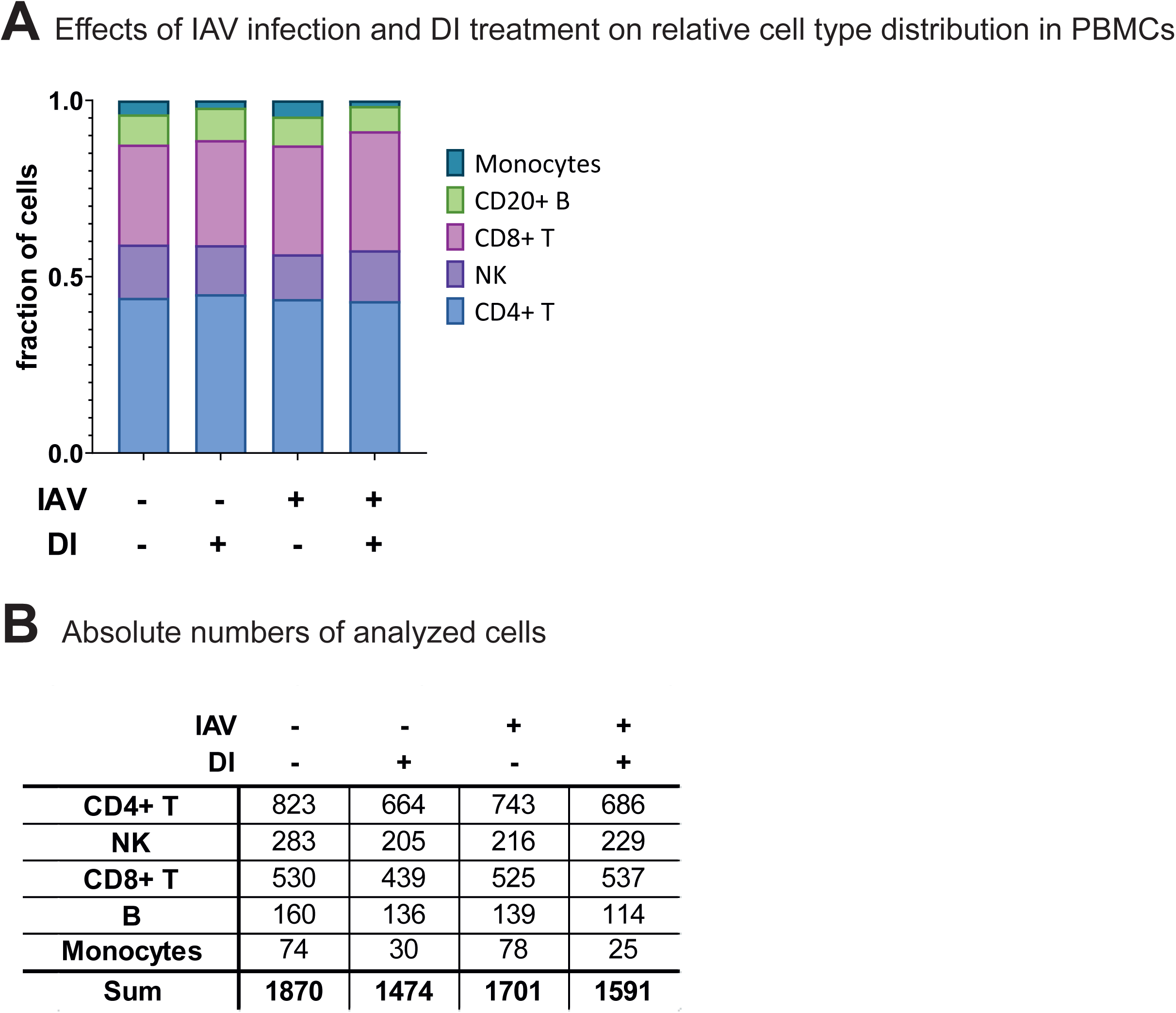
Effects of IAV infection and DI treatment on relative cell type distribution in PBMC. A reduction of CD14+ monocytes is apparent under DI treatment of both uninfected and infected PBMC. **A.** Percentages. **B.** Absolute numbers.

**Figure S18.**
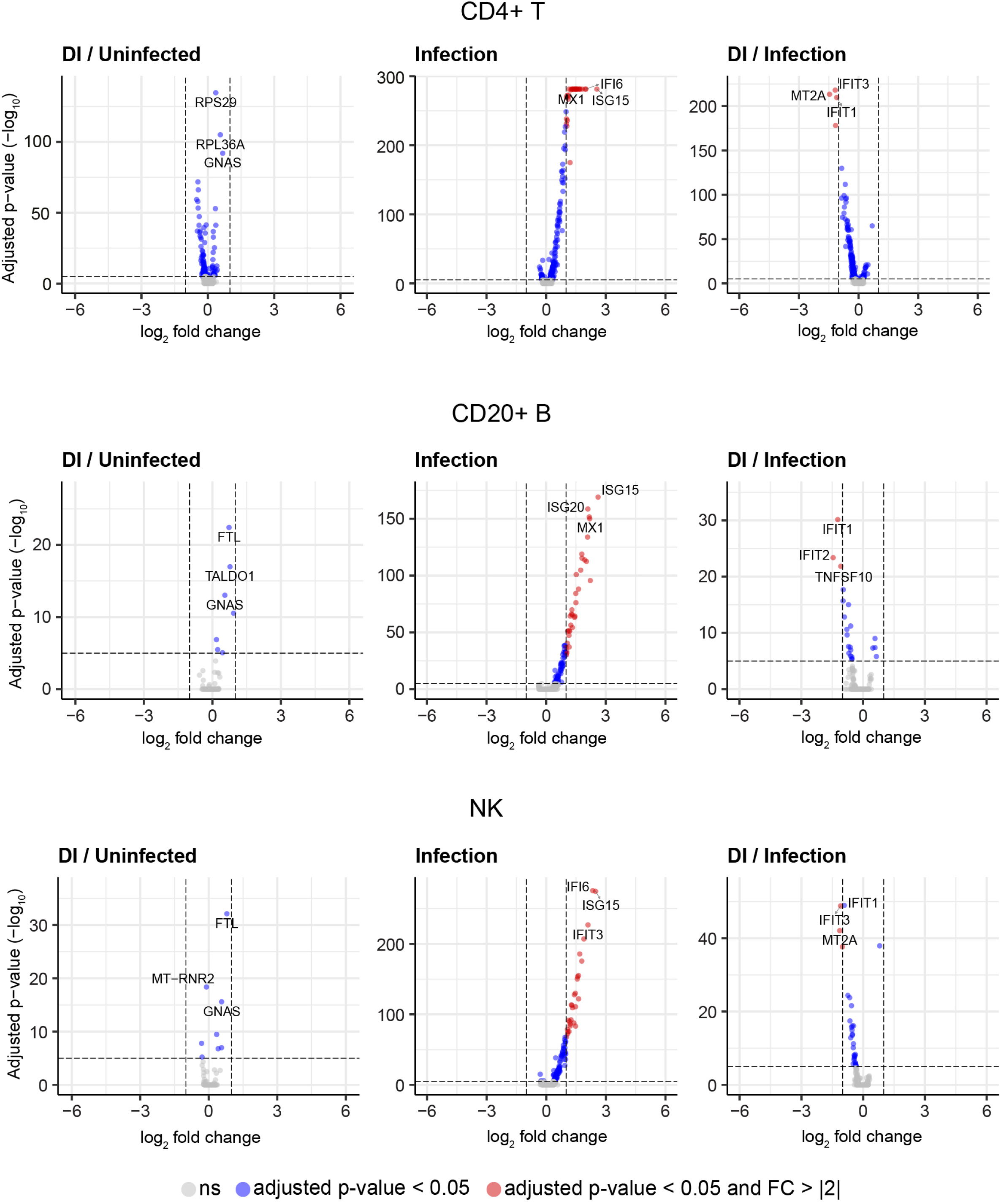
Effects of IAV infection and DI treatment on transcriptomes in the PBMC cell types not shown in Fig. 9. Analysis based on scRNAseq. Volcano plots comparing effects of DI treatment, with and without IAV infection, on CD4+ cells, NK cells, and B cells.

**Figure S19.**
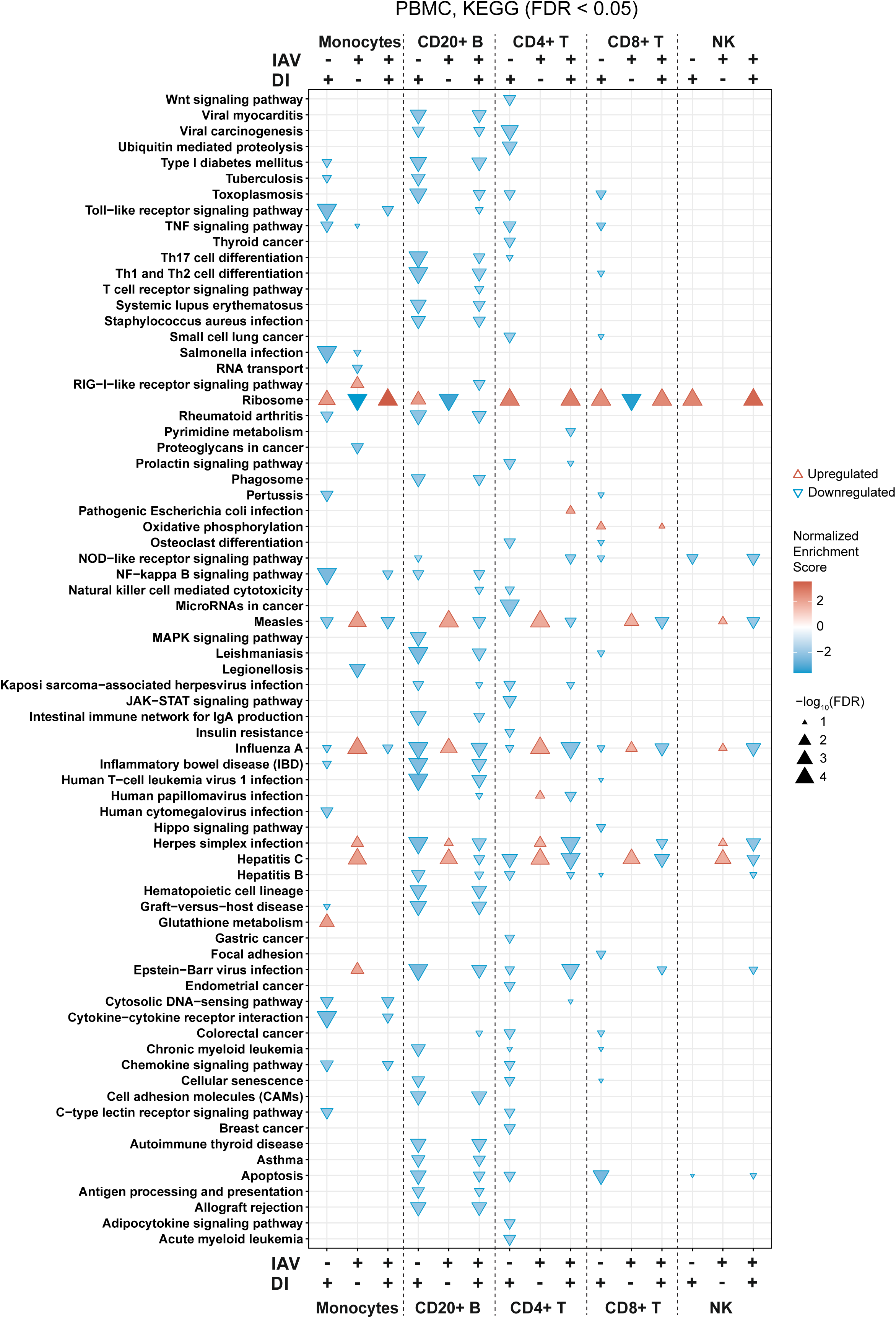
Cell-type specific KEGG pathway analysis of effects of DI on infected and uninfected monocytes, CD4 T cells, CD8 T cells, B cells, and NK cells. Analyses based on the scRNAseq data used for Fig. 9 and 10. Pathways that are enriched (FDR <0.05) in at least one cell type are shown.

**Figure S20.**
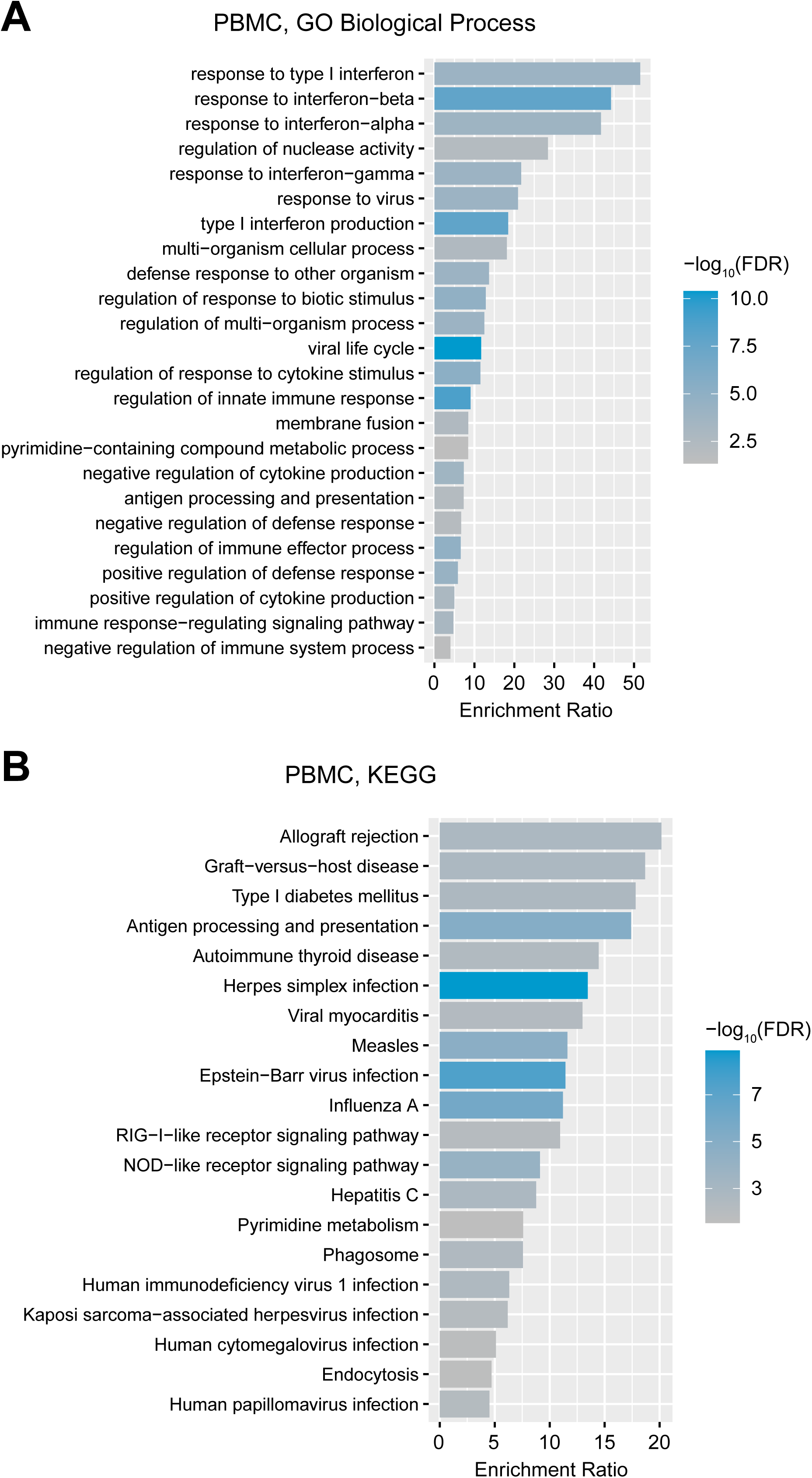
Identification of central pathways commonly regulated in monocytes, CD4 and CD8 T cells, B cells, and NK cells in response to IAV infection and DI treatment. GO Biological Process (A) and KEGG (B) functional enrichment analyses of all genes commonly differentially expressed in monocytes, CD4 and CD8 T cells, B cells, and NK cells in response to IAV infection and DI treatment (center of Venn diagram shown in Fig. 9H, based on scRNAseq analysis of PBMC).

